# Patterns of presence-absence variation of NLRs across populations of *Solanum chilense* are clade-dependent and mainly shaped by past demographic history

**DOI:** 10.1101/2023.10.13.562278

**Authors:** Gustavo A. Silva-Arias, Edeline Gagnon, Surya Hembrom, Alexander Fastner, Muhammad Ramzan Khan, Remco Stam, Aurélien Tellier

## Abstract

Understanding the evolution of pathogen resistance genes (nucleotide-binding site-leucine-rich repeats, also known as NLRs) within a species requires a comprehensive examination of factors that affect gene loss and gain. We present a new reference genome of *Solanum chilense*, that leads to an increased number and more accurate annotation of NLRs. Next, using a target-capture approach, we quantify the presence-absence variation (PAV) of NLR *loci* across 20 populations from different habitats. We build a rigorous pipeline to validate the identification of PAV of NLRs, then show that PAV is larger within populations than between populations, suggesting that maintenance of NLR diversity is linked to population dynamics. Furthermore, the amount of PAV is not correlated with the NLR presence in gene clusters in the genome, but rather with the past demographic history of the species, with loss of NLRs in diverging populations at the distribution edges and smaller population sizes. Finally, using a redundancy analysis, we find limited evidence of PAV being linked to environmental gradients. Our results contradict the classic assumptions of the important selective role of PAV for NLRs, and suggest that NLRs PAV is driven by random processes (and weak selection) in an outcrossing plant with high nucleotide diversity.

## Introduction

With reduced costs for high-throughput sequencing technologies, it becomes possible to quantify the genetic diversity of large sets of individuals across populations and test theoretical predictions regarding the importance of gene families (copy number variation, gene presence/absence) versus single gene processes in plant local adaptation to their biotic and abiotic environment. This amounts to quantifying the respective contributions of neutral random evolutionary forces (mutation, recombination, genetic drift, demography and spatial structure) versus local selective forces (positive, balancing or purifying selection) within and between populations using whole genome SNP polymorphism and gene presence-absence variation (PAV). Based on the population genetics literature on selection at single or multiple loci (reviewed in e.g. Stephan, 2016; Barghi *et al*., 2020; Johri *et al*., 2022) and the evolution of gene families by gene duplication/deletion (reviewed in e.g. Innan & Kondrashov, 2010; Kondrashov, 2012; Birchler & Yang, 2022), two complementary scenarios of adaptive evolution can be suggested and tested.

First, adaptation can occur via selected mutations at one (or several) single copy gene(s). Plant local adaptation to biotic or abiotic stress happens by selective sweep or balancing selection at such mutations, which would differ between populations. Such mutations, and the causal genes, exhibit higher population differentiation than the genomic average (identified for example using *F*_ST_ scans; Savolainen *et al*., 2013; Hoban *et al*., 2016) and can be confirmed by a significant association with specific environmental variables (using redundancy analysis, RDA, e.g. Capblancq & Forester, 2021; Wei *et al*., 2023a). *F*_ST_ scans and RDA analyses should ensure to disentangle the selection footprints from the neutral population past demography and spatial structure (e.g. Capblancq *et al*., 2018) as population mutation rate of the species determines the rate at which new mutations appear and the strength of genetic drift (Savolainen *et al*., 2013; Hoban *et al*., 2016).

In the second scenario, adaptation relies on PAV, the various gene copies being collectively subjected to (positive or balancing) local selection (e.g. varying across different habitats). In the case of neo-subfunctionalization, purifying selection maintains the original gene function at one copy, while other ones may evolve a new advantageous function by mutation (e.g. Ohta, 1976; Kondrashov, 2012). Selection may, instead or in addition, occur concomitantly on the number of gene copies itself, for example when nucleotide diversity across all gene copies is advantageous but too many copies can become disadvantageous (Innan & Kondrashov, 2010; Otto *et al*., 2022 and references therein). Here, PAV is likely more important than nucleotide diversity to generate novel adaptive features. We would expect that under local adaptation, PAV would exhibit higher differentiation between populations (measured for example as *F*_ST_ at the PAV level) and selected PAV to be correlated with environmental variables (as assessed by RDA). Then, the two main determinants of the speed of PAV driven local adaptation are (in contrast to scenario one) 1) the population recombination rate of the species which determines the rate at which new haplotypes appear, and 2) the effective size which determines the strength of genetic drift (Innan & Kondrashov, 2010; Otto *et al*., 2022).

We are interested in Nucleotide-binding site-leucine-rich repeats genes (known as NLRs or NB-LRR), which are vital components of disease resistance in all plant families (Gao *et al*., 2018; Barragan & Weigel, 2021). The diversity of these loci is intensely studied because of the need to enhance plant resistance against pathogens, but remains challenging. The study of NLRs is complicated by their repetitive nature and high variation in copy numbers (from single to hundreds of copies) across the genome in different plant species (Wersch & Li, 2019; Barragan & Weigel, 2021). NLR are functionally diverse and part of complex gene signalling networks. NLR genes called “helpers” are highly connected (hubs), while other so-called “sensors” are more peripheral (Wu *et al*., 2017a). The evolvability of NLR loci thus depends on their function, redundancy, and interactions within the network (Wu *et al*., 2017a; Stam *et al*., 2019b).

The extent to which diverse evolutionary mechanisms shape NLR gene evolution remains a topic of debate (Stahl *et al*., 1999; Bergelson *et al*., 2001; Holub, 2001; Märkle *et al*., 2022). Interspecific comparison of NLR sequences suggest that there is rapid evolutionary change of NLR families both as presence-absence of paralogs and at the nucleotide level (Gao *et al*., 2018; Liu *et al*., 2021), explained as a birth-and-death (turn-over) process of gene duplication, natural selection and deletion of unnecessary (or even deleterious) paralogs (Michelmore & Meyers, 1998), yielding the definition of pan-NLRome at the inter-specific level (Seong *et al*., 2020; Li *et al*., 2023b). However, in order to quantify the speed of NLR turn-over, that is namely the role of the underpinning random genomic processes, it is necessary to study NLR diversity at the intra-specific level. A main determinant of the NLR evolutionary trajectory is the effective population size of populations and species, which determines the duplication/deletion and gene conversion rates (Hörger *et al*., 2012) and the strength of selection (coevolution with pathogens) versus genetic drift and spatial structure (Tellier *et al*., 2014). In species with large effective population sizes (e.g. outcrossing) and large effective mutation rate, adaptation at NLRs could occur by mutation at single genes and to a lesser extent by PAV variation. However, if selection occurs on PAV as well, it should also be efficient and detectable (as both population mutation and recombination rates are high). On the other hand, in species with small population mutation rate (e.g. selfing and/or suffering recent bottlenecks) but with high population recombination rate (due to high per site recombination), adaptation may occur predominantly by selection on PAV. The process of NLR gene duplication and family expansion/contraction would also be affected by neutral processes. There is thus a need to disentangle neutral from selective effects at the PAV distribution and/or the polymorphism (SNPs) level (e.g. Stam *et al*., 2019b).

Studies in the model system *Arabidopsis thaliana* and crops report high rates of intra-specific presence-absence variation (PAV) as well as clustering of NLRs in the genome (Wersch & Li, 2019; Barragan & Weigel, 2021). Performing target capture on 64 *Arabidopsis* accessions, van der Weyer *et al*. (2019) confirmed levels of PAV which ranged from 167 to 251 loci per accession, with only half being present in most accessions. Such extensive PAV led to the notion of a pan-NLRome defining the core set of loci shared between individuals. Thus, PAV in NLR clades is assumed to result from a dynamic process of gene duplication/deletion generating different homeologs on which local selection can act (birth and death process; Michelmore & Meyers, 1998), and would underpin adaptation of populations to different pathogen pressures (van der Weyer *et al*., 2019; Barragan & Weigel, 2021). However, proof is lacking regarding the role of selection (positive and/or balancing; (Stahl *et al*., 1999; Bergelson *et al*., 2001; Hörger *et al*., 2012; Tellier *et al*., 2014) in promoting neo-subfunctionalization, and/or driving presence-absence diversity.

Early studies in *A. thaliana* (Bergelson *et al*., 2001; Tian *et al*., 2002; Allen *et al*., 2004; Bakker *et al*., 2006; Huard-Chauveau *et al*., 2013) find modest evidence for pervasive positive or balancing selection on polymorphisms at NLR loci, as confirmed by later genome scans for footprints of selection (Wu *et al*., 2017b). Van de Weyer et al. (2019) show a correlation in population genetics statistics between sensor and helper gene pairs. However, such an effect may also be due to linkage disequilibrium and variable rates of recombination along the genome. Lee & Chae (2020) took these analyses further and showed that there are differences in PAV across NLR clades and that radiations (clade expansion) of NLRs were not common. It remains unclear whether PAV between populations of *A. thaliana* across different environments can be attributed solely to neutral effects of 1) demographic events of population bottleneck and post-glacial age expansion, and/or 2) the relatively small local effective population sizes due to high selfing rate and absence of seed banks (Alonso-Blanco *et al*., 2016; Sellinger *et al*., 2020). Furthermore, studies investigating NLR diversity and evolution in other wild species are rare and often limited to few loci (e.g. Rose *et al*., 2007, 2011).

To fill this gap, we study *Solanum chilense*, an obligate outcrossing species which presents strong patterns of local adaptation with known past demographic events of colonisation of various habitats around the Atacama desert (Böndel *et al*., 2015; Stam *et al*., 2019b; Wei *et al*., 2023a). It exhibits high local effective population sizes, due to the seed banks (Tellier *et al*., 2011) and mild bottlenecks of colonisation, and thus high genetic diversity at the SNP level. Selection occurring at genes for abiotic adaptation has been documented at SNP (Mboup *et al*., 2012; Böndel *et al*., 2015; Wei *et al*., 2023a) and PAV levels (Fischer *et al*., 2011; Wei *et al*., 2023b). We previously found a few NLRs under positive selection for local adaptation based on SNPs (Stam *et al*., 2019b), on a subset recovered through a PoolSeq method and a draft genome assembly of *S. chilense* (Stam et al., 2019a).

Here we assess the occurrence of selection pressure for PAV vs. SNP diversity. We first complete a new genome reference of *S. chilense* with chromosome-size scaffolds and annotate NLR genes. We then sequence by target capture this set of NLRs across 200 plants of *S. chilense* from 20 populations to study PAV and SNP diversity, focusing on conserved NB-ARC domains. To examine if selection is occurring we 1) examine and compare Fst values for PAV vs. SNP, and 2) test for selection of PAV and SNP data through an RDA analysis using representative environmental variables. Contrary to SNP diversity which shows a footprint for selection, the main determinants of PAV within and between populations do not appear to be related to selection, but rather to their putative molecular functions in the genome as well as their past demographic (neutral) population history.

## Material and methods

### Genome scaffolding, annotation and visualisation

High-molecular weight DNA was sent to Dovetail Genomics (Santa Cruz, CA, USA) to construct Chicago libraries. The libraries were sequenced on an Illumina HiSeqX with 150 bp paired-end reads. Using the draft assembly as input (Stam *et al*., 2019a), the HiRise scaffolding pipeline, to build super scaffolds, was done at Dovetail Genomics (proprietary protocol). Short-read sequences generated from Chicago and HiRise libraries are available at ENA (PRJNA508893). A full description of the methods and bioinformatic pipelines used to polish, map and assess the quality of the genome, as well as annotation methods, are available in Supporting Information Methods S1.

### Annotation of NLR loci in the reference genome

The NLR-annotator pipeline (Steuernagel *et al*., 2020) was used to identify NLR loci in the new reference genome, using default commands, resulting in the identification of 278 NLR loci. We next focused exclusively on loci containing complete nucleotide-binding site domain-encoded genes (NB-ARC), as *de novo* assembly of the LRR domains of NLRs can be complicated without long-read sequencing and manual curation (Jupe *et al*., 2013; Witek *et al*., 2021; Barragan & Weigel, 2021). The NB-ARC domain has a high conservation across the plant kingdom (Shao *et al*., 2016) and can be confidently resolved (Barragan & Weigel, 2021). This approach has been used successfully in previous NLR studies (Lee & Chae, 2020; Safdari *et al*., 2021). A total of 177 NB-ARC loci were identified, of which seven were eliminated because they represented duplicated or overlapping loci (Dataset S2). A full account of our approach to link and annotate these genes based on phylogenetic reconstructions, which allows us to compare with the previous G3 genome (Stam et al. 2019), is found in Supporting Information Methods S2.

### Evaluation of NB-ARC presence-absence variation (PAV) and allelic diversity (SNPs)

#### Sampling

We sampled 200 plants from 38 accessions and grew each plant in individual pots under standard glasshouse conditions at the GHL Dürnast plant research facilities of the TUM School of Life Sciences. The sampling includes 18 accessions (hereafter populations) from the central region (10 samples each) arranged in four central valleys (CV) representing replicates of elevation gradients (Fig. 3a). The remaining 20 plants were sampled from scattered populations (one plant per locality) from the two divergent lineages (south-highland SH, and south-coast SC). Plants were grown from individual seeds obtained from the Tomato Genetics Resource Center (TGRC, University of California, Davis, USA; <u>tgrc.ucdavis.edu</u>). Genomic DNA was extracted using the DNAeasy extraction kit from Qiagen following the instructions of the supplier.

#### Target enrichment and sequencing

We designed 1,959 probes targeting coding sequences of 170 NB-ARC regions annotated in this study. Probe synthesis, library preparation, and sequencing were carried out at RAPiD Genomics LLC (Gainesville, CA) using SureSelectXT (Agilent Technologies, CA) enrichment system followed by Illumina Hiseq 2000 sequencing to generate paired-end 150-bp reads.

#### Assembly of NB-ARC loci, PAV detection and validation

We used the Captus pipeline (Ortiz *et al*., 2023) that automatizes the processes of raw read quality check and trimming, *de novo* assembly, and extraction of the NB-ARC target loci from the full set assembled contigs for each sample. Reference loci for the extraction process corresponded to the 170 NB-ARC domains previously identified with NLR-Annotator. Briefly, the Captus pipeline uses Scipio to identify gene structures given a protein sequence. Captus also identifies intron-exon borders and splice sites, and can cope with loci assembled over multiple contigs (fragmented assemblies). We performed the extraction with a relaxed set of parameters (minimum Scipio score = 0.13, minimum identity percentage to reference proteins 65%, and minimum coverage percentage of reference protein to consider a hit by a contig = 20%).

The identification of PAV was further refined with the identification of cut-off values of identity and coverage statistics, which are described in detail in Supporting Information Methods S3. The resulting high-confidence (HC) dataset consisted of a total of 156 NB-ARC loci sequenced for 186 sequenced individuals across 20 populations (Dataset S3).

We scored PAV by: (1) assigning a binary character for each locus, namely a locus that either did not show any presence-absence variation across the entire dataset, or showed absence for at least one individual; (2) assigning a quantitative variable per locus indicating the locus frequency (presence) across a set of sequences. We calculated both quantities for several sets of sequences: across all 186 individuals, within each of the 20 populations sampled, as well as within the six geographic regions. We tested our approach on the RENseq long-read data from (Seong *et al*., 2020) using our set of 156 NB-ARC loci annotated in *S. chilense* revealing consistent patterns of PAV as our interspecific data set, but showing more absences in more variable NB-ARC clusters according to the divergence of the included species (Fig. S6). We find 3,074 NB-ARC loci/sample combinations (10.6% of the total full data set; 156 loci * 186 plants) showing multiple hits above the identity and coverage thresholds (Fig. S4) indicating possible copy number variation (CNV) of the locus in each sample (darker colours in Fig. S1). This CNV signal is consistent with higher mapping rates than expected based on the read mapping to the reference (see Fig. S5). All of these loci also scored positive for PAV across the data set (as method (1)), thus these loci were treated as regular loci with PAV. An *F*_ST_ analogous statistic to quantify the PAV pairwise population differentiation was calculated using custom R scripts and visualised using boxplots and heatmap with ggplot in R.

#### Short-read alignment, variant calling and diversity of NB-ARC loci

We mapped the filtered read data to the reference genome using bwa-mem version 0.7.17 (Li & Durbin, 2009; Li, 2013) with default settings. We then sorted, fix mates, mark duplicates and retain only concordantly mapped reads, output BAM files using the samtools version 1.10 (Li *et al*., 2009) and calculated per locus depth of coverage using mosdepth version 0.3.3 (Pedersen & Quinlan, 2018). We then created an “all sites” vcf file containing both invariant and variant sites using the “mpileup” and “call” methods implemented in BCFtools version 1.19 (Danecek *et al*., 2021). Detected variants were filtered to retain bi-allelic SNPs with a site quality >30 and a depth of coverage greater than 5x and smaller than 200x using vcftools version 0.1.16 (Danecek *et al*., 2011).

Pairwise population genetic differentiation (*F*_ST_) for each locus was calculated using pixy version 1.2.7 (Korunes & Samuk, 2021) for each NB-ARC region, specifying the start and end position of each locus in a bed file. We also calculated per population allele frequencies for each locus using vcftools. All the scripts used for these calculations are available in the project repository (https://gitlab.lrz.de/population_genetics/Schilense_newref).

### Attributes of NB-ARC loci

For each of the loci with complete NB-ARC domain, we scored them for attributes which possibly influence their PAV across populations:

(1) The location of NLRs as physical clusters, as defined by Andolfo *et al*. (2021). Genes within a maximum distance of 200,000 bp apart were considered clustered. We calculated the number of clusters based on all NLRs, including loci with incomplete NB-ARC domains (278 loci).
(2) The putative function of the NLR as being either helper, sensor, or non-sensor genes (*sensu* Wu *et al*., 2017). We defined helper genes as loci that were sister to the core four core NRC sequences previously identified in *A. thaliana* and tomato species (Wu *et al*., 2017a; Stam *et al*., 2019b,a) leading to an expanded clade of 10 loci (Fig 1b, Dataset S2; hereafter referred to as “Helper+”).
(3) The annotation of NLRs as TNL, CNL or RNL, based on the annotation method using phylogenies, described above.

**Figure 1.**
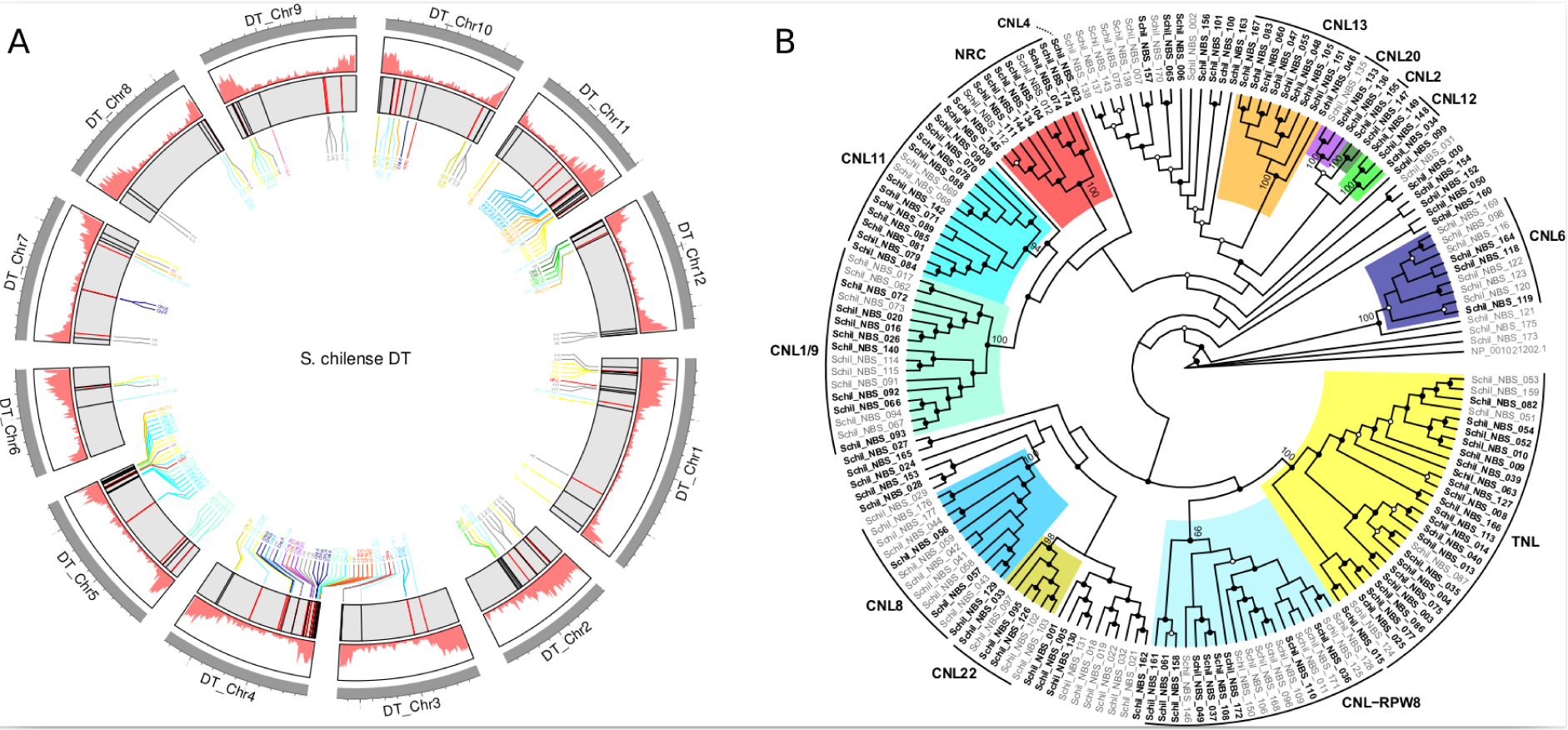
**(a) Genome assembly of *S. chilense.*** (1) Density of mRNA genes; (2) NLRs found in the *S. chilense genome* are shown, loci in black were found in the previous version of the genome (Stam *et al*., 2019a), whereas loci in red are new to this version; (3) annotation labels indicating the clades to which the Nb-Arc loci correspond (coloured as in Fig. 1B). **(b) Phylogeny of the NLR domains**. Maximum Likelihood phylogeny of the NB-ARC domains recovered from the Dovetail genome assembly in *S. chilense*, using NLR-Annotator. Clades are coloured using the same colour scheme as used in Stam *et al*. (2019a). Nodes with UltraFastBootstrap support of 95% and above are indicated with black circles, whereas bootstrap support above between 80% and 94% are indicated with white circles. Nodes without circles have branch support values below 80%. For each of the 15 NLR clades, we indicated the UltraFastBootstrap value next to the crown node of each respective clade. NB-ARC tip names in bold represent sequences that formed a strongly supported sister pair (95% Bootstrap support and above) with NB-ARC sequences identified in Stam *et al*. (2019a).

### Visualisation and chi-square tests

To visualise PAV across individuals and the 20 populations sampled, barplots, boxplots and UpSet plots were produced, using the following R packages: ggplot2, dplyr, stringr, tidyverse, ggtext, and UpSetR (Wickham, 2016, 2022; Gehlenborg, 2019; Wickham *et al*., 2019, 2022; Wilke & Wiernik, 2022). In addition, we carried out χ2-tests (chisq.test, as implemented in the “stats” package in R) to determine whether there were any significant differences in the frequency of various attributes.

### Comparison of pairwise F*_ST_* values for PAV and SNPs

Pairwise *F*_ST_ values for PAV of all NB-ARC loci included in this study was calculated using custom R scripts (https://gitlab.lrz.de/population_genetics/Schilense_newref). Variation of *F*_ST_ values for different NB-ARC loci across different populations were visualised using boxplots and heatmaps, and we used Wilcoxon signed rank tests and the Spearman rank correlation, to determine if we could detect any significant differences or correlations between *F*_ST_ of PAV and SNP diversity.

### Environmental correlation analysis with PAV and SNPs

#### Environmental data and occurrence records

We tested for correlation between PAV of NLR loci and five environmental variables reflecting strong constraints for growth and survival of *S. chilense* and the dataset on SNPs of the same NB-ARC loci. The selected variables included a proxy for cold stress (minimum temperatures of the coldest month, bio6), temperature seasonality (temperature annual range, bio7), seasonality of rainfall (precipitation amount of the wettest month, bio13), intra-annual variability of cloud frequency (MODCF_intraannualSD), and mean cloud frequency of September (MODCF_monthlymean_09), which is the month with the strongest coastal fogs in the Lomas ecosystems (Ruhm *et al*., 2022). Temperature and precipitation variables were derived from CHELSA climate data, whereas cloud data was taken from the Global 1-km Cloud Cover dataset (Wilson & Jetz, 2016) downloaded from the EarthEnv portal (Amatulli *et al*., 2018). All maps were at a 30 arcsec spatial resolution (ca. 1 km).

Occurrence records of the 38 localities present in our samples were downloaded from the TGRC and double-checked. Correlations among variables were verified and a standard PCA of the environmental data was carried out, using custom R scripts. Finally, we took the average value of all localities of the Southern Highland Group and the Southern Coastal Group, considering each group as a single population. This was justified given the environmental distinctiveness of these localities based on our preliminary PCA results.

#### Environmental analyses

We ran an initial PCA analysis of the frequency of the 156 NB-ARC loci across the 20 populations, and tested for association between frequency of specific alleles to environmental variables using a redundancy analysis (*rda* function from the vegan package in R; Oksanen *et al*., 2022). Prior to running the analysis, the NLR frequency data (PAV) was transformed using the Hellinger distance. Both the SNP and PAV data were modelled across the 20 localities as a function of the environmental variables. The significance of RDA-constrained axes was assessed using the *anova.cca* function. Finally, we ran a partial RDA, with the four central valleys (CV1 to CV4), the SH and SC groups as covariable factors using the entire 156 NB-ARC dataset or a sub-set of genes with the 113 NB-ARC loci from the CNL clade. The same analyses were carried out on the population allelic diversity measured on the same 156 NB-ARC loci.

## Results

### Dovetail scaffolding resolves the *Solanum chilense* genome to near chromosome level and identified additional NLRs

The assembly for the new reference genome of *S. chilense* has 12 chromosome-level scaffolds (110.22 to 53.75 Mb) and 12,626 unplaced scaffolds, resulting in an L50/N50 of 6/71.61Mb. The BUSCO scores (based on *Solanales_odb10*) are 95% complete (93% single copy and 2% duplicated), 1.1% fragmented and 3.9% missing BUSCO genes, supporting the improved quality and completeness compared to other assemblies in *S. chilense* and other species belonging to the tomato clade (Table 1). We annotate 40,113 protein-coding features, of which 74% show InterPro functional annotations (Dataset S1).

**Table 1.**
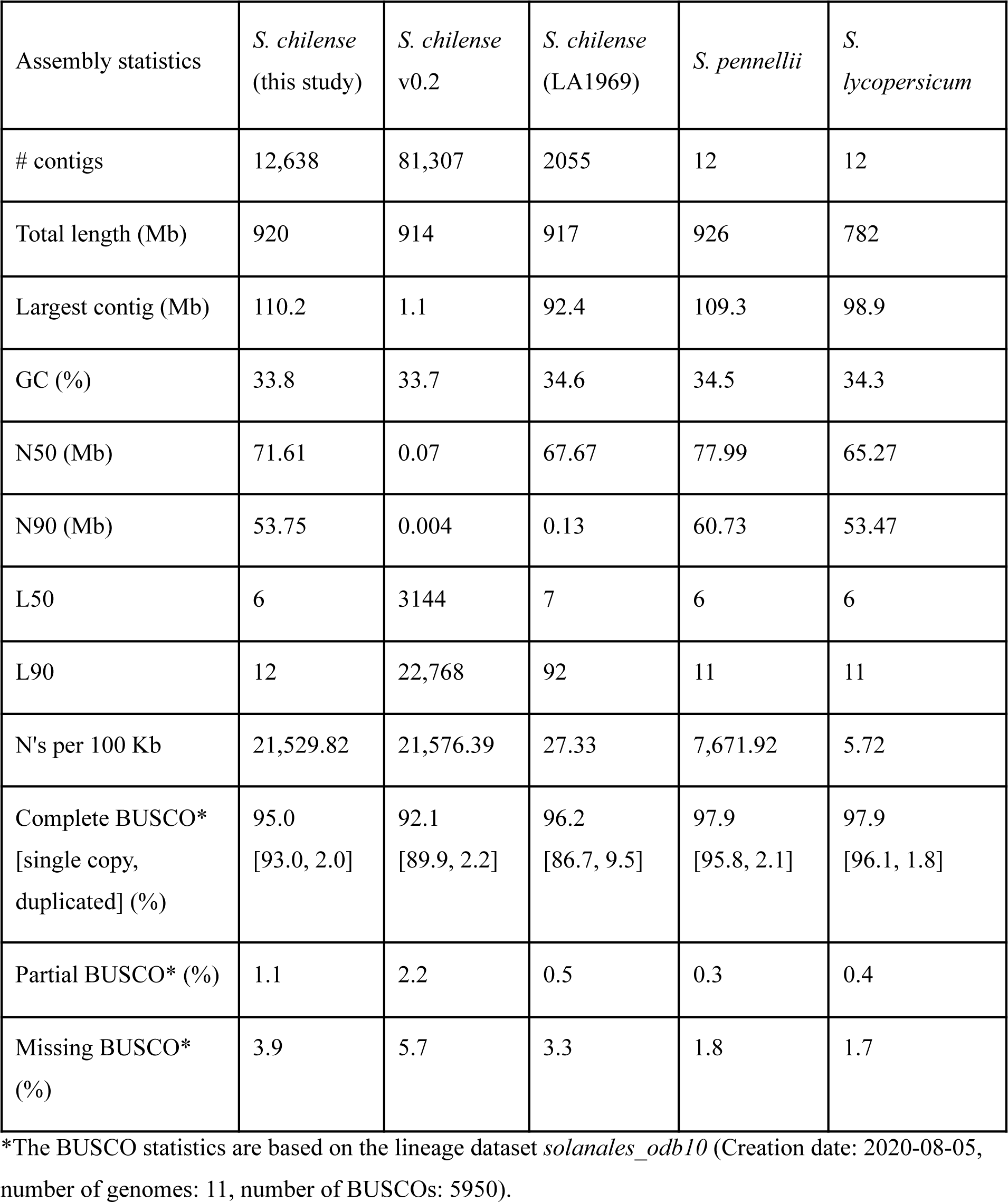
Assembly statistics of the new version of *Solanum chilense* reference and comparison with the previous assembly (Stam *et al*., 2019a), *S. chilense* (LA1969; Li et al., 2023a), *S. pennellii* (Bolger *et al*., 2014) and *S. lycopersicum* (Hosmani *et al*., 2019).

We identify 170 NLR sequences with complete NB-ARC domains (Fig. 1a, b), which is higher than the 134 NB-ARC sequences annotated in the previous genome assembly (Stam et al. 2019b), among which 123 (72%) NB-ARCs belong to the CNL clade, 27 and 20 genes belonging to the TNL and RNL clades, respectively. For six of the clades (CNL4, CNL12, CNL13, CNL21, CNL20, and NRC), we recover the same number of sequences in the new genome assembly (Table S1). We find variation in NLR numbers for nine other clades compared to the previous assembly (Table S1; (Stam *et al*., 2019a). In addition, we find another 15 novel NB-ARC sequences which could not be assigned to a clade (individual NLR: “ind”).

### Physical clustering of NLR loci and composition of clusters

When assessing the number of physical clusters of our NLR loci, we find 104 loci (61%) occuring in 57 clusters, and the top 20 clusters contain a third of all NLRs (97/278, 34%). Most clustered NB-ARCs are CNLs (78%, or 82/104 loci), 15 are TNLs, and 7 are RNLs. Cluster size ranges from 2 to 10 loci (median =2). Ten clusters are composed of a single NLR type (CNL or TNL), six of those are transcribed in the same direction and likely originated from tandem duplication events. We speculate that the remaining clusters originated from more complex genomic recombination events, as 13 have mixed compositions of different NB-ARC loci (including partial sequences; Dataset S2).

### NB-ARC loci show considerable PAV in S. chilense

Across the 186 individuals ([HC dataset; see methods]), we document an average of 137 NB-ARC loci (median=137, SD=3.96; max: 145, min: 123) per individual (Fig. 2a). 30 NB-ARC loci (20%) are fixed in terms of presence across all individuals and populations (15 TNLs, 14 CNLs, and one RNL). A total of 13 of these genes are singletons, whereas 17 are in clusters. The remaining 126 unfixed loci are absent in at least one sample, with 12 loci present in less than half of all the individuals sampled (Fig. 2b). Unlike in *A. thaliana*, we do not find a typical separation of core vs. cloud NLRs (van der Weyer *et al*., 2019), but rather a geometric abundance distribution with very few NLRs occurring less frequently (Fig. 2b). CNLs show higher PAV than TNLs. CNLs are on average present in 160 individuals (median=180, SD=39.3; max:186, min:18) and TNLs in 182, with a smaller variance (median=181.6, SD=10.5; max: 186, min:143). Differences in PAV frequencies are detected amongst different CNL clades: the CNL1/9 clade shows the greatest amount of variation, contrasting strongly for example with CNL-RPW8 (also known as RNLs; Fig. 2c). An overview of PAV for each NLR in each individual can be found in Fig. S1.

**Figure 2.**
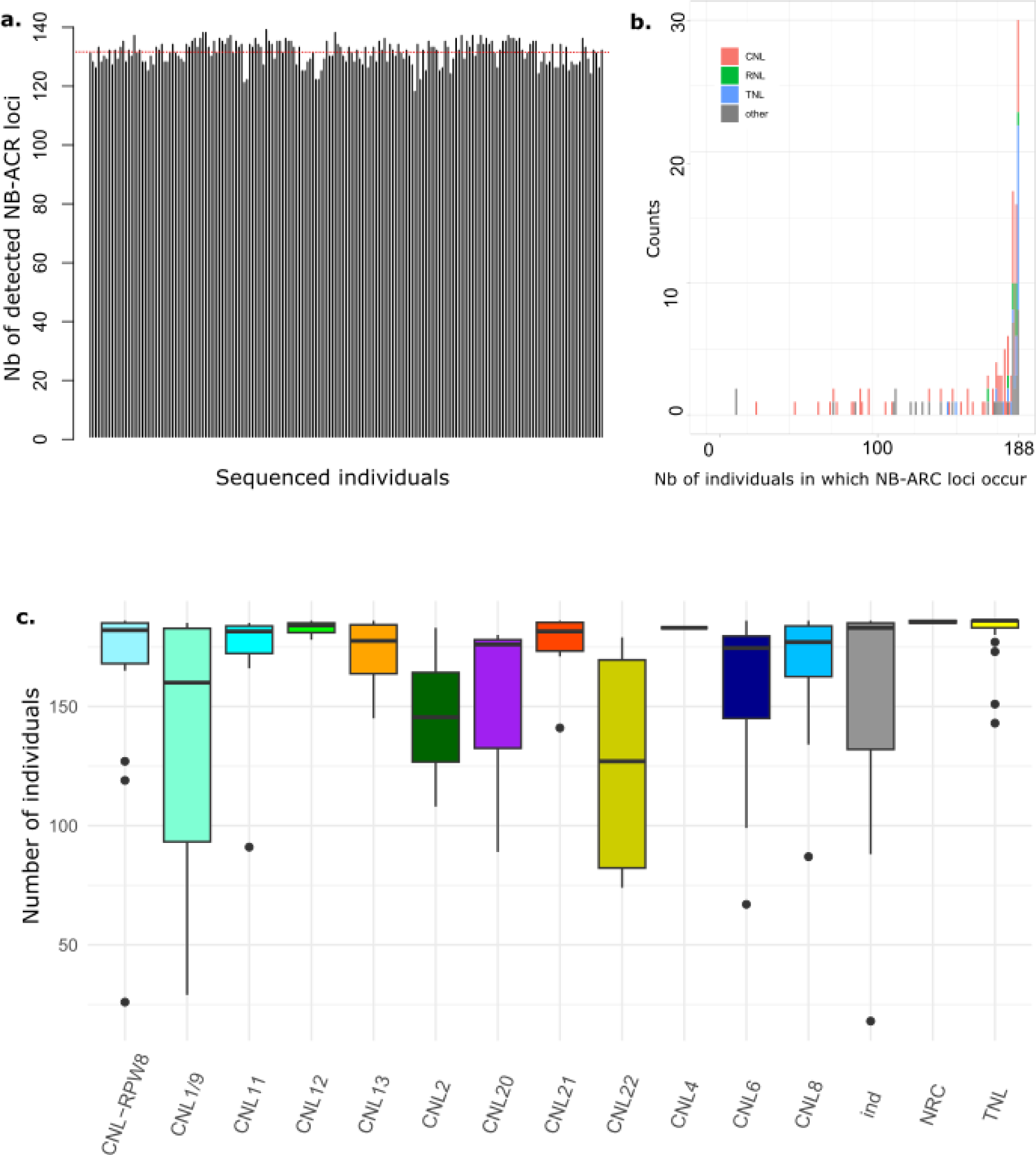
PAV in NB-ARC loci within *S. chilense.* (a) Total number of detected NB-ARC containing loci (y axis) for each of the sequenced individuals, each bar on the x axis represents a single individual; (b) Histogram showing the frequency distribution (y axis) of NB-ARC loci through our data set, shown as the number of individuals in which a locus occurs (x axis); (c) Barplot of NB-ARC genes per population, with each barplot showing the number of loci belonging to different NB-ARC clusters.

We find significant differences in the frequencies of loci with PAV based on NLR attributes for CNL vs. TNL loci, helper vs. sensor loci, and for sensor vs non-sensor loci, but not when comparing loci that are clustered vs. singleton (Table 2).

**Table 2.** Chi-square test results, comparing various attributes of NB-ARC loci. P-values below the threshold of 0.5 are indicated with an asterisk (*).

| **Comparison** | **X-square test** | **P-value** |
| --- | --- | --- |
| **Cnl/Tnl vs. PAV** | X-squared = 24.981, df = 1 | p-value = 5.789e-07* |
| **Cnl/Tnl vs. Clustered/Singleton** | X-squared = 0.410, df = 1 | p-value = 0.5222 |
| **PAV vs. Clustered/Singleton** | X-squared = 0.320 df=1 | p-value = 0.5715 |
| **Helper+/Sensor vs. PAV** | X-squared = 7.511, df = 1 | p-value = 0.006131* |
| **Helper+/Sensor vs. Clustered/Singleton** | X-squared = 2.205, df = 1 | p-value = 0.1375 |
| **Sensor/Non-Sensor vs. PAV** | X-squared = 7.045, df = 1 | p-value = 0.007951* |
| **Sensor/Non-Sensor vs. Clustered/Singleton** | X-squared = 0.057, df = 1 | p-value = 0.8109 |

### NB-ARC loci have high variation within population, yet are generally maintained within geographical groups

When recalculating the PAV of NB-ARC loci across our 20 populations (Fig. 3a) (loci considered as present if found in one individual of a population), we find that variation amongst populations is lower than compared to variation across all individuals. We identify an average of 153 NB-ARC loci (median=154, SD=2.68; max: 156, min: 145) per population (Fig. 3b), a total of 133 loci (85%) being found across all populations. From the remaining loci, 22 are found in most populations (12 to 19) and a single locus is found in just eight populations. Looking at PAV within these populations, we observe variation in the mean number of recovered NB-ARC loci per population of 130.9 to 140.6 loci, with small SD (2.1-5). The lowest average and median are in the SC population (mean=130.9, median=131, SD: 3.89) (Fig. 3c).

**Figure 3.**
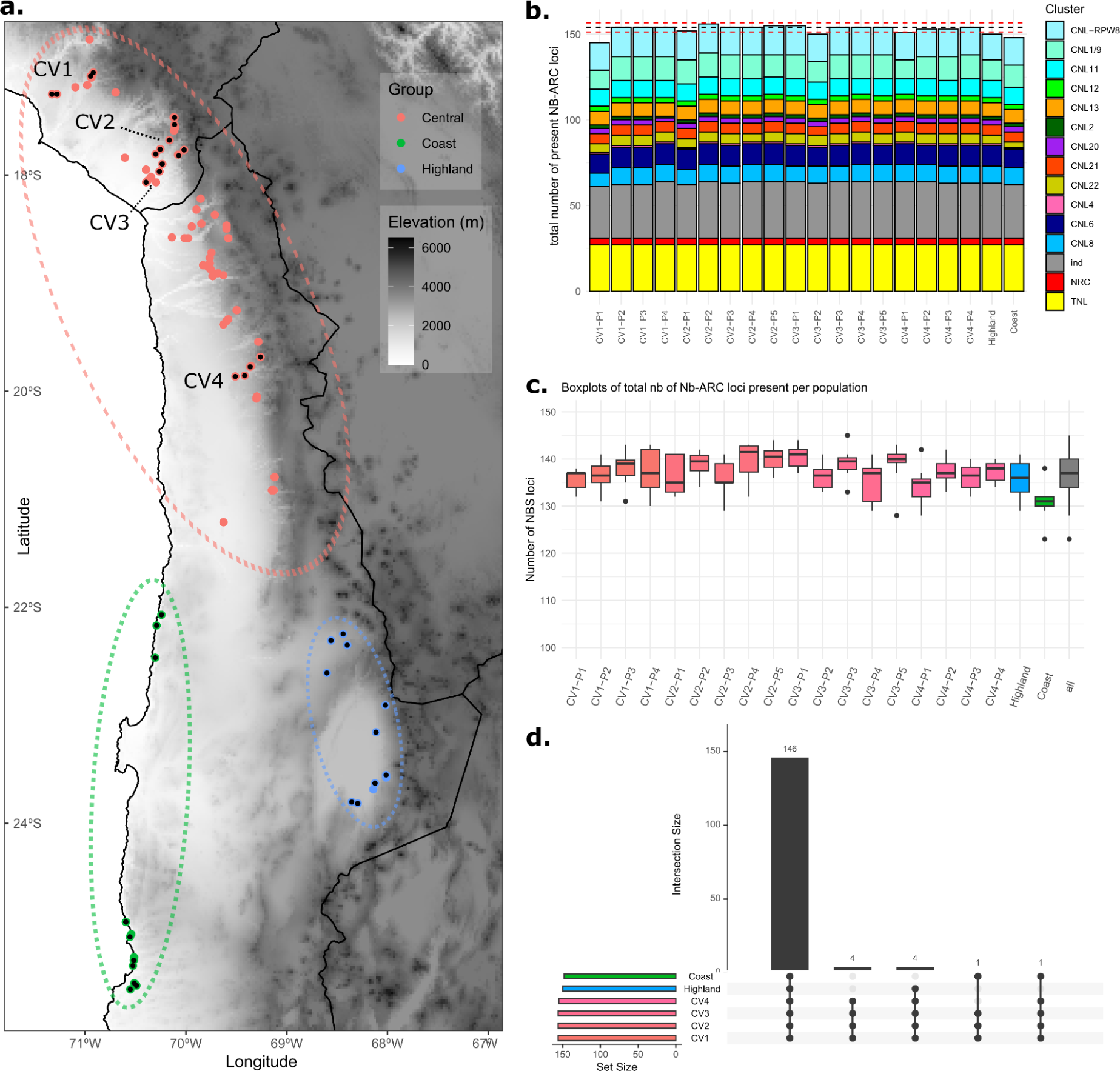
PAV of NB-ARC loci on a geographical scale. (a) Map of the localities of the TGRC samples that were sampled in this study, from the Central Valley, Southern Highland and Southern Coastal regions; colored dots represent known, active collection, the one with black dots are included in our study; (b) Barplot of NLR genes per population, with each barplot showing the number of loci belonging to different NB-ARC clusters; the solid blue line represents the median of NB-ARC loci across all populations, and the two dotted red lines represent the standard deviation interval; (c) Boxplots of the total number of NB-ARC loci per population; colours correspond to CV, SH and SC regions as shown in Fig 3A; (d) Upset plot of NB-ARC loci present across the six metapopulations of *S. chilense*, showing how many loci are shared between the six metapopulations, from the Central Valleys 1 to 4 (CV1 to CV4), the Southern Highland region (SHG), and Southern Coastal Group (SCG).

Next, when examining PAV for the six geographically defined regions included in this study, four valleys in the central region (CV1, CV2, CV3 and CV4) and two regions from the southern range (SC and SH; Fig. 3a), we find that all 156 loci are at least present in one or more individuals in the four CV regions, with the exception of one locus lost in CV4 and SH (Fig. 3d). Ten loci are not found in the SH and SC regions (Fig. 3d), and four loci are lost only in the SC region. One locus is lost only in the SH region, and four loci are lost shared by the SC and SH regions (Fig. 3d). All these loci are CNLs, the majority from the CNL1/9 and CNL22 categories. Most of these genes are not sensor or helper NLRs (8 out of 10) and are found physically clustered in the genome (6 out of 10).

### Comparison of F*_ST_* values based on nucleotide diversity and PAV

We observe lower values for population differentiation (*F*_ST_) for the PAV of NB-ARC loci compared to the nucleotide (SNP) diversity in the populations sampled in this study (Fig. S7). This suggests that mutation rather than gene duplication and deletion processes predominantly drive local differentiation at NB-ARC loci in *S. chilense*. The calculated *F*_ST_ values reveal a significant difference (Wilcoxon signed rank test; p-value = 3.05 × 10^−8^) showing greater average differentiation in nucleotide variation (mean: 0.2; sd: 0.08; min: 0.06 [CV4-P1 vs. CV4-P3]; max: 0.47 [CV1-P2 vs. Coast]) (Fig. S7b) than in locus PAV (mean: 0.17; sd: 0.05; min: 0.06 [CV4-P1 vs. CV4-P3]; max: 0.36 [CV1-P1 vs. CV3-P2]) (Fig. S7b) and confirmed by the distribution of pairwise *F*_ST_ values for all sampled populations (Fig. S7c). With the SNP data we recover the expected higher *F*_ST_ values for population pairs involving comparisons between the Central Regions (CV) and the Southern Highland and Coast populations (Fig. 3a). This pattern is very subtle in the pairwise *F*_ST_ obtained with the PAV (Fig. S7c). The SNP and PAV average *F*_ST_ are correlated (Spearman’s rank correlation rho=0.6; p-value = 2.2 × 10^−16^) supporting the idea that both are mainly shaped by demographic processes.

The per-locus pairwise *F*_ST_ values for PAV exhibit no significant differences between the CNL, TNL and RNL loci (Kruskal-Wallis test; p-value = 0.6; Fig. S8a). Slightly higher values are found for CNL loci (mean: 0.13; sd: 0.16; min: 0 [183 pairwise observations]; max: 1 [40 pairwise observations]), compared to RNL (mean: 0.12; sd: 0.16; min: 0 [27 pairwise observations]; max: 1 [6 pairwise observations]), and lower values are found for TNL (mean: 0.12; sd: 0.14; min: 0 [11 pairwise observations]; max: 0.65 [13 pairwise observations]) (Fig. S8a). On the other hand, the per-locus pairwise *F*_ST_ values for SNP data show significant differences among NB-ARC classes (Kruskal-Wallis test; p-value = 1.13 × 10^−94^; Fig. S8b). CNL loci also have larger *F*_ST_ (mean: 0.2; sd: 0.15; min: 0 [226 pairwise observations]; max: 0.97 [CV3-P2 vs. Coast]), compared to RNL (mean: 0.16; sd: 0.13; min: 0 [14 pairwise observations]; max: 0.98 [CV3-P2 vs. Highland]), and lower values are found for TNL (mean: 0.22; sd: 0.14; min: 0 [26 pairwise observations]; max: 0.82 [CV4-P4 vs. Coast]) (Fig. S8b).

### Contrasting results for RDA of NB-ARC SNPs, PAV and environmental variables

The PCA of environmental variables (Fig. 4a) shows CV1, CV2 and CV3 tend to cluster and are ordered along an altitudinal gradient. The most southern Central group (CV4) forms its own cluster, as do SC and SH, confirming that they differ in their environment. In the PCA of NB-ARC frequencies across 20 populations (Fig. 4b), we find three main clusters: one including populations of CV1, one with populations of CV4, and a third containing nearly all populations from CV2 and CV3. Coastal and Highland populations, along with a single population from CV3, occupy a distinct position in the PCA scatterplot. These results suggest that there is strong population structure of NB-ARC frequencies, which needs to be taken into account when considering whether PAV of NB-ARC might be correlated with environmental variables.

**Figure 4.**
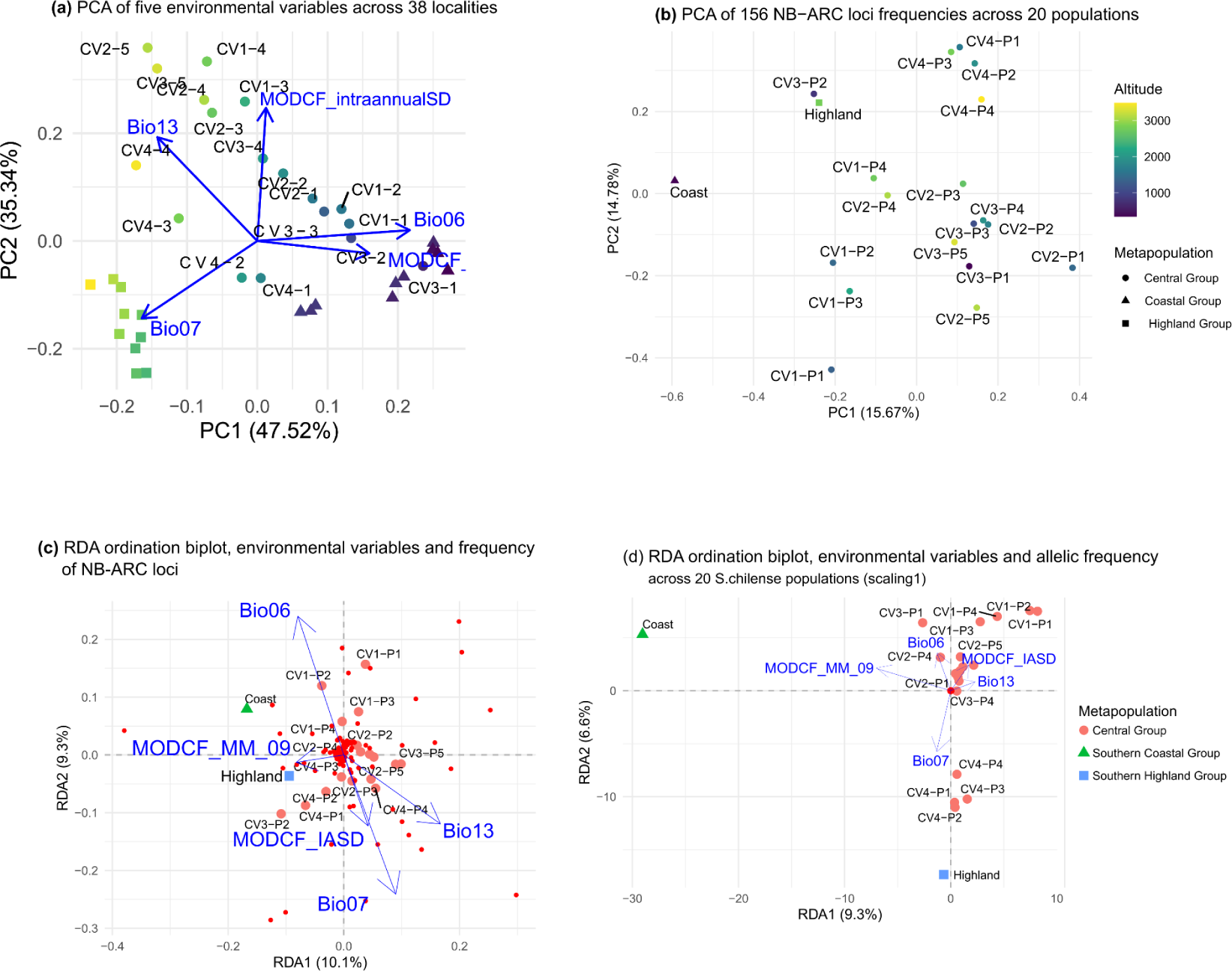
**(a) PCA based on the five environmental variables, across 38 localities of *S. chilense* present in this study.** The three main groups are represented by different shapes (circle: Coastal Group; triangle: Southern Coastal Group; square: Southern Highland group), and the dots are coloured according to elevation; (b) PCA of the PAV (frequency) of the 156 NB-ARC loci across 20 populations; (c) RDA analysis of PAV of NB-ARC loci, scaling 1; (d) RDA based on SNP allele frequency of NB-ARC loci, scaling 1.

The RDA with PAV data is not significant in the ANOVA permutation test, for both the dataset containing all 156 NB-ARC loci (p=0.184, Fig. 4c), and the second analysis focuses on CNL loci (p=0.218, figure not shown). This result remains unchanged when accounting for population structure with the partial RDA analysis (ANOVA permutation test: p=0.09, figure not shown).

In contrast, the RDA using the SNPs of the 156 NB-ARC loci is found to be significant when using the ANOVA permutation tests (p=0.002, fig 4d) with the first axis of the RDA (p= 0.06, Fig. 4d). A clear distinction between populations is observed in the RDA biplots, similar to what is found in the PCA of environmental variables: the RDA 1 clearly helps to distinguish the Coastal population, whereas the RDA 2 helps distinguish a cluster of CV1, CV2, and CV3, a cluster of CV4, and the highland group (Fig. 4d). A partial RDA is also performed taking into account population structure of the Central Valley Groups, but the ANOVA permutation test is not significant (p=0.6, figure not shown). Taken together, we interpret these results as suggesting that there is some weak evidence for association of particular NB-ARC alleles with environmental variables.

## Discussion

The new reference genome of *S. chilense* and refined annotation of coding regions allows us to identify 278 NLRs, of which 170 are full length with NB-ARC domain. Previous reports in wild tomatoes range between 204 and 265 across six self-compatible wild tomato species and the self-incompatible *S. habrochaites* (Seong *et al*., 2020, 2022). We annotate about 18% more sensor NLRs than previously (Table S1). The genes for which we detect the largest increase in the number of genes are non-sensor loci (33% more, Table S1), suggesting that these genes are subject to even less constraint than the genes involved in NLR networks.

Like previous studies, we find PAV at NLRs between individuals of the same species (van der Weyer *et al*., 2019; Seong *et al*., 2020; Lee & Chae, 2020), with only 30 NB-ARC loci fixed across all individuals. Within and between populations the level of conservation of genes remained relatively high in comparison, and have similar patterns to other wild relatives of tomatoes, for example more PAV in CNLs vs. TNL loci (Seong *et al*., 2020). Furthermore, the pattern of PAV is rather consistent with the past demographic history of the species. Loss of genes occurs chiefly in Southern Coastal and Highland populations, that are derived from two colonisation events to novel habitats and characterised by mild bottlenecks (Böndel *et al*., 2015; Stam *et al*., 2019b; Wei *et al*., 2023a). However, contrary to previous studies (van der Weyer *et al*., 2019; Lee & Chae, 2020), we do not find that NB-ARC loci clustered in the genome have significant differences in PAV compared to singleton NB-ARC loci (Table 2). This result suggests that the rates of duplication and deletion events in *S. chilense* are irrespective of their arrangement in clusters along the genome, and the absence of NLR “hotspots” of diversification.

With regards to evidence for selection, our study shows a lack of correlation of PAV of NLR loci with environmental variables based on the RDA (Fig 4). Evidence from early studies finding cluster size differences between different ecotypes provide limited evidence for selection on NLR loci with regard to environment variables (Botella *et al*., 1998; Noël *et al*., 1999). To our knowledge, PAV in NLR genes according to an altitudinal gradient has only been reported once at the *CHS3* and *CSAI* NLR genes in *A. thaliana* fixed in high-altitude populations (Günther *et al*., 2016). As both genes also play a role in the response to different environmental cues, such as chilling stress, such environmental correlations may be very rare events. Lee & Chae (2020) investigated the cluster diversity and size of NLR based on the *A. thaliana* panNLRome and have not found any correlation between cluster size or cluster size expansion with altitude, latitude or longitude. We conclude that PAV at NLRs in *S. chilense* (and *A. thaliana*) are likely not driven in a simple manner by presence-absence of different pathogens across the range of habitats (and environmental gradient) of the species.

In contrast, weak evidence is observed for SNP association of particular NB-ARC loci with environmental variables, as confirmed by the *F_ST_*comparison between SNPs and PAV. This weak footprint for selection at SNP level was also found by Stam et al. (2019). We conclude that genetic diversity on which selection acts is based on point mutations (and indels) in *S. chilense,* rather than on PAV, confirming our hypotheses in the introduction regarding the genomic bases for adaptation in outcrossing plant species with high effective population sizes. In *S. chilense,* the effective population size is on the same order of magnitude as the population recombination rate (Tellier *et al*., 2011; Wei *et al*., 2023a), and novel DNA mutations are as frequent as recombination events to generate novel variants. As a result, partial mildly deleterious gene copies may then be strongly counter-selected. If PAV would be advantageous in generating novel haplotypes (by mutation, gene conversion or homologous recombination), positive or balancing selection at PAV underpinning local adaptation should be detectable (see the example of *Rcr3* in the sister species *S. peruvianum*; Hörger *et al*., 2012).

Our results do not negate that PAV may be an important source of variability for NLRs, especially for species with small effective population size such as *A. thaliana* or crops (due to high selfing, fragmented populations and history of bottleneck and colonisation of habitats). As outlined in our introduction, it is still unknown to which extent PAV is driven by neutral or selective processes, and we attempt in this paper to provide a framework to test this hypothesis. In *A. thaliana*, the population recombination rate is five times higher than the population mutation rates (Nordborg, 2000; Sellinger *et al*., 2020), and this ratio changed during the transition from outcrossing to selfing (Strütt *et al*., 2023). Homologous recombination and gene conversion would thus be more efficient at creating novel NLRs variants (haplotypes) than DNA mutation. On one hand, such unequal crossing over may generate additional non-functional (or partial) NLR paralogs that can be observed as PAV. On the other hand, large numbers of NLRs could hinder the integrity of the genome (by generating hybrid necrosis; Chae *et al*., 2014). Thus, if purifying selection against these extra-numerous, possibly deleterious, NLR variants is relatively weak and/or genetic drift is strong due to a low effective population size, partial gene copies can be maintained or even reach fixation. Consequently, we speculate that the large PAV and hot-spot effect of clustering in *A. thaliana*, may be in large part due to neutral processes driven by small local effective population sizes, until the respective contributions of (weak) selective and demographic processes can be disentangled.

Variation between individuals and maintenance of diversity at NLR loci across *S. chilense* species range could be due to balancing selection by pathogens or to seedbanking. These are hypothesised to be particularly important due to El Niño climatic fluctuations which leads to long-term cyclical rain and population dynamics, and to intermittent gene flow and dispersal of seeds within the different valleys (Tellier *et al*., 2011). Neutral demographic processes are also likely to play a role in shaping PAV and SNP variation at NLRs. This stems from the weak selection pressure for the plant to adapt to pathogens in arid habitats, as indicated by the large and variable resistance response to infection by various pathogens (Stam *et al*., 2017; Kahlon *et al*., 2020, 2023). Our RDA considers environmental variables, such as humidity, temperature and rainfall, which are classically linked with the presence/absence of plant pathogens, but we cannot exclude that other variables may be more relevant and would correlate with PAV across habitats. In *S. chilense*, (putative) natural pathogens have been identified, but more work is needed to quantify their respective pressure across different populations (Stam *et al*., 2017; Kahlon *et al*., 2020; Schmey *et al*., 2023). The lack of correlation between NLR and environmental variables could also be due to the redundancy of gene function in the NLR network (Wu *et al*., 2017a) affecting the efficiency of natural selection for gene duplication/loss.

Our approach has both strengths and limitations with regards to the study of NLR genes. First, the Illumina-based target capture approach allows us to estimate loss and absence of loci with regards to our reference genome. However, our results are slightly biassed as we cannot accurately estimate the gain of particular loci. Yet, losing a gene is still more likely than gaining one. Considering the rigour of our probe design, we are confident that even with PacBio sequencing, we would not have necessarily found significantly more gains in loci than losses. Our results and interpretations are thus likely not affected by this slight bias. Second, germplasm can bias population genomic studies, due to the possibility of ex-situ genetic drift caused by the method of propagating and maintaining collections. This issue was discussed in previous studies (Wei *et al*., 2023a,b), and was taken into account here by screening out samples for outlier populations with peculiar low levels of nucleotide diversity. The two populations with lowest NLR diversity that were removed were the oldest collected germplasm accessions multiplied several times at TGRC (see Böndel *et al*., 2015). As our results only marginally rely on accurate estimations of allele frequencies, our results are likely robust to this effect.

In conclusion, we propose that combining high-quality reference genome with rigorous assembly and screening of NLR diversity via target capture is especially relevant for studying NLR loci diversity at the population genomic level in crops and non-model species. Our hypotheses may help to understand the observation of a low number of NLRs and low PAV in several species. Several lineages have undergone substantial contractions of total number of NLRs (Baggs *et al*., 2020), and a more recent study examining NLR diversity and evolution across 300 angiosperm genomes found that this contraction seemed to occur in plant lineages undergoing ecological specialisation related to aquatic, parasitic and carnivorous growth forms (Liu *et al*., 2021). These dynamics are likely to vary according to different evolutionary and ecological contexts, and the complexity and redundancy of the NLR/pathogen interaction network within each species (Adachi & Kamoun, 2022). For example, in *Amborella*, weak PAV of genes related to biotic stress was observed, in sharp contrast with abiotic stress genes (Hu *et al*., 2022), presumably through lack of pathogen pressure and limited distribution in remote oceanic islands, not unlike *S. chilense*. Expanding these studies to more species is crucial for understanding how habitat loss and fragmentation can impact plant health and resistance in the wild.

## Acknowledgements

We thank Edgardo Ortiz for advice in the processes with Captus. EG acknowledges support by the European Union’s Horizon 2020 research and innovation program under the Marie Skłodowska-Curie grant agreement no. 899987. AT acknowledges funding from DFG (Deutscheforschungsgemeinschaft) grant no.: 317616126 (TE809/7-1). AT and MRK were supported by a DAAD Germany - Pakistan exchange grant; GAS-A was funded by the Technical University of Munich. RS acknowledges DFG grant no.: 170483403 (SFB924). We thank the GHL Dürnast (TUM) for the use of plant growing facilities, and the Tomato Genetics Resource Center (TGRC) of the University of California, Davis for generously providing us with the seeds of the *S. chilense* accessions.

## Author contributions

GS, AT, EG and RS planned and conceptualised the research; EG, GS, AF, SH carried out the analyses in the paper; AT, MRK and RS obtained funding; EG, GS, AT, RS all contributed significantly to the writing of the manuscript, with all co-authors reviewing and approving the final version.

## Data availability

The data that support the findings of this study are openly available in the European Nucleotide Archive at https://www.ebi.ac.uk/ena with the following accession numbers. SRA ERR11268525 (Chicago sequencing), SRA ERR11268526 (Hi-C sequencing) and BioProject PRJEB61272 (Target sequencing data). Reference genome sequence and annotation files are available on the Solgenomics website (https://solgenomics.net/ftp/genomes/Solanum_chilense/Gustavo/). The code (and Dataset S4) for implementing the analyses used in this paper can be found on our GitLab repository: https://gitlab.lrz.de/population_genetics/Schilense_newref.

## Supporting Information

**Figure S1**. Presence/Absence variation and copy number variation matrix of NB-ARC loci of *Solanum chilense* detected through *de novo* assembly approach across 186 samples included in the sequence capture data set, whole-genome data from the reference genome sample (LA3111_t13; used to assemble the reference genome in Stam *et al*., 2019a), and 30 whole-genome sequencing samples from Wei *et al*. (2023a). Some loci show higher mapping rates than expected based on the mapping rates in other loci, indicating possible CNV of the locus in these accessions. These loci are displayed in a darker colour.

**Figure S2**. Dotplot representation of sequence alignment of the new assembly of *Solanum chilense* and *S. pennellii*.

**Figure S3**. Maximum Likelihood phylogeny of the NB-ARC domains recovered from the Dovetail and the short-read sequence assembly from *S. chilense*, using NLR-Annotator. Clades are coloured according to the same scheme as used in Stam *et al*. (2019a). Nodes with UltraFastBootstrap support of 95% and above are indicated with white circles, whereas branch support of 80% to 94% are indicated with black circles. Numbers next to the left of the nodes indicate ultrafast bootstrap support, although these are not represented for nodes with support of 95% and above. Tip names in red represent NB-ARC sequences identified in Stam *et al*. (2019a).

**Figure S4**. 2D density plot showing the distribution of Identity percentage and Coverage percentage of all assembled contigs to the 170 NB-ARC reference protein sequences annotated in the new reference. Dashed lines indicate the threshold values chosen to filter the extraction.

**Figure S5**. Distribution of depth of coverage for each NB-ARC locus annotated in the new reference genome of *Solanum chilense* measured after read alignment of the 200 samples from the targeted sequencing (left panels) and 30 samples from whole-genome sequencing (right panels).

**Figure S6** NB-ARC loci presence-absence variation matrix obtained with long-read RENseq data from Seong *et al*. (2020) including *S. lycopersicum*, five wild tomatoes, and two additional (non-*Solanum*) Solanaceae species.

**Figure S7** Average population differentiation *F*_ST_ estimated on NB-ARC presence-absence variation (PAV, left panel) and nucleotide variation (SNP, right panel).

**Figure S8** Distribution of per-locus NB-ARC population pairwise differentiation *F*_ST_ estimated on presence-absence variation (PAV, top panel) and nucleotide variation (SNP, bottom panel).

**Table S1. Comparison of the 170 NB-ARC-containing NLRs recovered in the new versus the previous version *S. chilense* genomes, for each NLR clade.** Numbers in parenthesis indicate the number of new sequences that were identified in the new genome reference sequence.

## Supplementary datasets

**Dataset S1.** Protein-coding features from the new reference genome.

**Dataset S2.** Predicted genes that were annotated and considered functional based on searches against the InterPro and UniRef90 protein signature databases.

**Dataset S3.** Presence Absence table for 186 individuals and 156 loci (indicate excluded pops here), with copy number variation detected for each loci.

**Dataset S4.** Alignments of complete NB-ARC domains of reference genome with previous Phylogenies (in the Gitlab repository).

