## Supplementary files, Figures and Tables for "Patterns of presence-absence variation of NLRs across populations of *Solanum chilense* are clade-dependent and mainly shaped by past demographic history"

### *New Phytologist* Supporting Information



#### Article acceptance date: T.B.D.

**The following Supporting Information is available for this article:**

**Methods S1 – *Genome scaffolding, annotation and visualisation***

**Methods S2 – *Annotation of NLR loci in the reference genome***

**Methods S1 -** ***Genome scaffolding, annotation and visualisation***

High-molecular weight DNA was sent to Dovetail Genomics (Santa Cruz, CA, USA) to construct Chicago libraries. The Chicago libraries were sequenced on an Illumina HiSeqX with 150 bp paired-end reads. Using the draft assembly as input (Stam *et al.*, 2019), the HiRise scaffolding pipeline, used to build super scaffolds, was done at Dovetail Genomics using their proprietary protocol. Short-read sequences generated from Chicago and HiRise libraries are available at ENA (PRJNA508893).

Using Pilon v1.23 (Walker *et al.*, 2014) we ran one round of polishing onto the 12 chromosome-size contigs. Using BWA v0.7.17 we first mapped the paired short-read sequences of the same individual (LA3111_t13) from (Stam *et al.*, 2019). The sorted alignments were then used as input in Pilon in full-correction mode. The quality of the genome assembly was evaluated in Quast v5.2.0 (Gurevich *et al.*, 2013) which calculated basic statistics such as total length, GC content, N50/90, L50/90 and the number and size of the contigs. We also evaluated the completeness of the assembly using BUSCO v5.4.5 (Seppey *et al.*, 2019) based on the database Solanales v10 (Kriventseva *et al.*, 2019); creation date: 2020-08-05, number of genomes: 11, number of BUSCOs: 5950). The assignment of chromosome-size scaffolds to each respective chromosome was done by aligning the 12 biggest scaffolds of the new reference to the genome sequence of *Solanum pennellii* (Bolger *et al.*, 2014), using the web application D-GENIES (Cabanettes & Klopp, 2018) and minimap v2 (Li, 2021) (Fig S2).

Before annotating the assembly, we soft-masked the repetitive sequences using the benchmarking pipeline EDTA (Ou *et al.*, 2019). The repeat analysis shows that 60.06% of the *S. chilense* genome comprise transposable elements, the majority being long terminal repeat (LTR) retroelements (36.1%), mostly Gypsy-LTRs (26.09%). We performed structural gene annotation using BRAKER v2.1.5 (Brůna *et al.*, 2021) which relies on GeneMark-ET (Lomsadze *et al.*, 2014) (Lomsadze *et al.*, 2014) and AUGUSTUS v3.2.3

(Stanke *et al.*, 2006). We first performed *ab initio* gene prediction. Subsequently, we performed 10 runs of evidence-based gene prediction in BRAKER using transcriptomic paired-end libraries from leaf tissues of six individuals of *S. chilense* LA3111 (PRJNA474106; Stam *et al.*, 2019). For that, we mapped the RNA reads to the reference for genome annotation using BBmap (Bushnell, 2014), after filtering the adapters and low-quality reads with the Captus pipeline (Ortiz *et al.*, 2023). Additionally, we used Liftoff v1.3.0 (Shumate & Salzberg, 2021) with options -a 0.9 -s 0.8 -copies, to transfer the existing gene model annotations from *S. chilense* (Stam *et al.*, 2019) and *S. pennellii* (GCF_001406875.1) into the new assembly. The three sets of predicted/transferred genes were then merged to generate a nonredundant reference gene set using EvidenceModeler v1.1.1 (Haas *et al.*, 2008). For functional annotations, Blastp v2.12.0+ (Camacho *et al.*, 2009) with a threshold E-value of <1e−5) was used to align the protein sequences to the UniRef90 database (The UniProt Consortium, 2021), and only the best-matched targets were retained. InterProScan v5.52-86.0 (Jones *et al.*, 2014) was used to annotate motifs and domains by searching against ProDom (Bru *et al.*, 2005), CDD, Gene3D, PRINTS, PFAM, SMART, PANTHER, SUPERFAMILY, TIGRFAM, and PROSITE databases. Gene Ontology (Harris *et al.*, 2005) annotations were obtained based on the InterPro entries.

**Method S2 -** ***Annotation of NLR loci in the reference genome***

The NLR-annotator pipeline (Steuernagel *et al.*, 2020) was used to identify NLR loci in the new reference genome, using default commands. Focusing solely on the protein alignments of the nucleotide-binding site domain-encoded genes (NB-ARC), we recovered a total of 177 loci, of which seven were eliminated because they represented duplicated or overlapping NLR loci (Dataset S2). The output from NLR-annotator was first used to reconstruct a phylogenetic tree based on the protein alignments of the NB-ARC domains of the identified 170 NLRs; the outgroup sequence used in NLR-annotator (NP_001021202.1, Cell Death Protein 4 CDP4), was also included in the phylogenetic analyses. Protein sequences were aligned using default settings in MAFFT v.7, through the online portal service (Nakamura *et al.*, 2018), with manual curation subsequently carried out in Jalview (Waterhouse *et al.*, 2009). A maximum likelihood tree of the NB-ARC domain alignment was done with IQ-Tree2 (Minh *et al.*, 2020), using ModelFinder to find the best substitution model for the alignment (in all cases the model JTT+F+R6 is selected according to the Bayesian Information Criterion). In addition, a total of 1000 UltraFastBootstrap were carried out to calculate branch support. Visualisation of the phylogeny was carried out in R and Inkscape, using the packages ggtree (Yu *et al.*, 2017), ggtreeExtra (Xu *et al.*, 2021) and ggplot2 (Wickham, 2016).

Previous NLR sequences from the Stam *et al.* (2019) genome assembly identified a total of 134 NB-ARC domains that can be grouped into 15 different clades. To understand whether the new reference genome was able to recover these same sequences, the 170 NB-ARC loci from the new reference were aligned with the NB-ARC loci from the previous assembly using MAFFT v.7. A maximum likelihood phylogeny was calculated and visualised, using the same methods as described above. The NB-ARC loci from the new reference were assigned to different NLR clades based on whether they formed solid sister pairs with annotated NB-ARC loci from the previous assembly (bootstrap support above 95%). For all other sequences, the assignment to NLR clades was based on a visual inspection of the phylogenetic relationships (Fig 1a and S3).

In order to visualise and compare the assemblies of the previous and new references, as well as the location of the NB-ARC domains along the genome, we used the package "circulize" in R (Gu *et al.*, 2014). We also visualise the density of mRNA genes annotated along each of the scaffolds. For the new reference genome, we recovered 12 major scaffolds (corresponding to 12 chromosomes), in which the 170 detected NB-ARC loci were all found (Fig. 1b). In comparison, the previous assembly had 134 NB-ARC loci across a subset of the genome composed of 113 scaffolds.

**Methods S3 – *Assembly of NB-ARC loci, PAV detection and validation***

We used the Captus pipeline (Ortiz *et al.*, 2023) that automatizes the processes of raw read quality check and trimming, *de novo* assembly, and extraction of the NB-ARC target loci from the full set assembled contigs for each sample. Reference loci for the extraction process corresponded to the 170 NB-ARC domains previously identified with NLR-Annotator. Briefly, the Captus pipeline uses Scipio to identify gene structures given a protein sequence. Captus also identifies intron-exon borders and splice sites, and can cope with loci assembled over multiple contigs (fragmented assemblies). We performed the extraction with a relaxed set of parameters (minimum Scipio score = 0.13, minimum identity percentage to reference proteins 65%, and minimum coverage percentage of reference protein to consider a hit by a contig = 20%).

The identification of PAV was further refined with the identification of cut-off values of identity and coverage statistics. To establish the cut-off values, the distribution of identity and coverage was evaluated for all extracted loci and visualised in 2D density plots with ggplot in R (Fig. S4). The distribution of the statistics for the complete data set showed that most of the assembled contigs have high coverage (> 95%). Within them, we found two clearly defined peaks of identity values. In order to avoid erroneous captures of paralogs we delimited a strict identity cutoff value of 94% which points to the group of assembled sequences of high confidence.

To compare the patterns of presence/absence variation (PAV) of NB-ARC loci identified with the Captus pipeline, we applied the above-described method to 1) the original ~130x short-read data used to assemble the first version of the genome of *S. chilense* (Stam *et al.*, 2019; LA3111_REFERENCE in Fig. S1), and 2) 30 whole genome sequencing samples (Wei *et al.*, 2023; marked with WGS suffixes in Fig. S1). The loci were annotated for the individual LA3111_REFERENCE. We found nine loci missing in the reference sample which are likely due to technical error during sequencing or assembly. These nine loci were thus removed from further analysis. Preliminary data exploration led to the exclusion of 14 samples from the target capture data set, which consistently showed a lower number of total NLR loci and lower read depth of coverage after short-read alignment (mapping) to the reference genome (see details of mapping procedure below). Furthermore, a total of 14 NB-ARC loci were also excluded, as they showed discordant PAV patterns between the target capture and WGS data or very low or no read coverage after short-read alignment (Fig. S5), likely because target capture failed to capture them due to the lack of ad hoc probes for these loci. Then we remove them to avoid over-interpretation of the data. The resulting high-confidence (HC) dataset consisted of a total of 156 NB-ARC loci sequenced for 186 sequenced individuals across 20 populations (Dataset S3).

New Phytologist

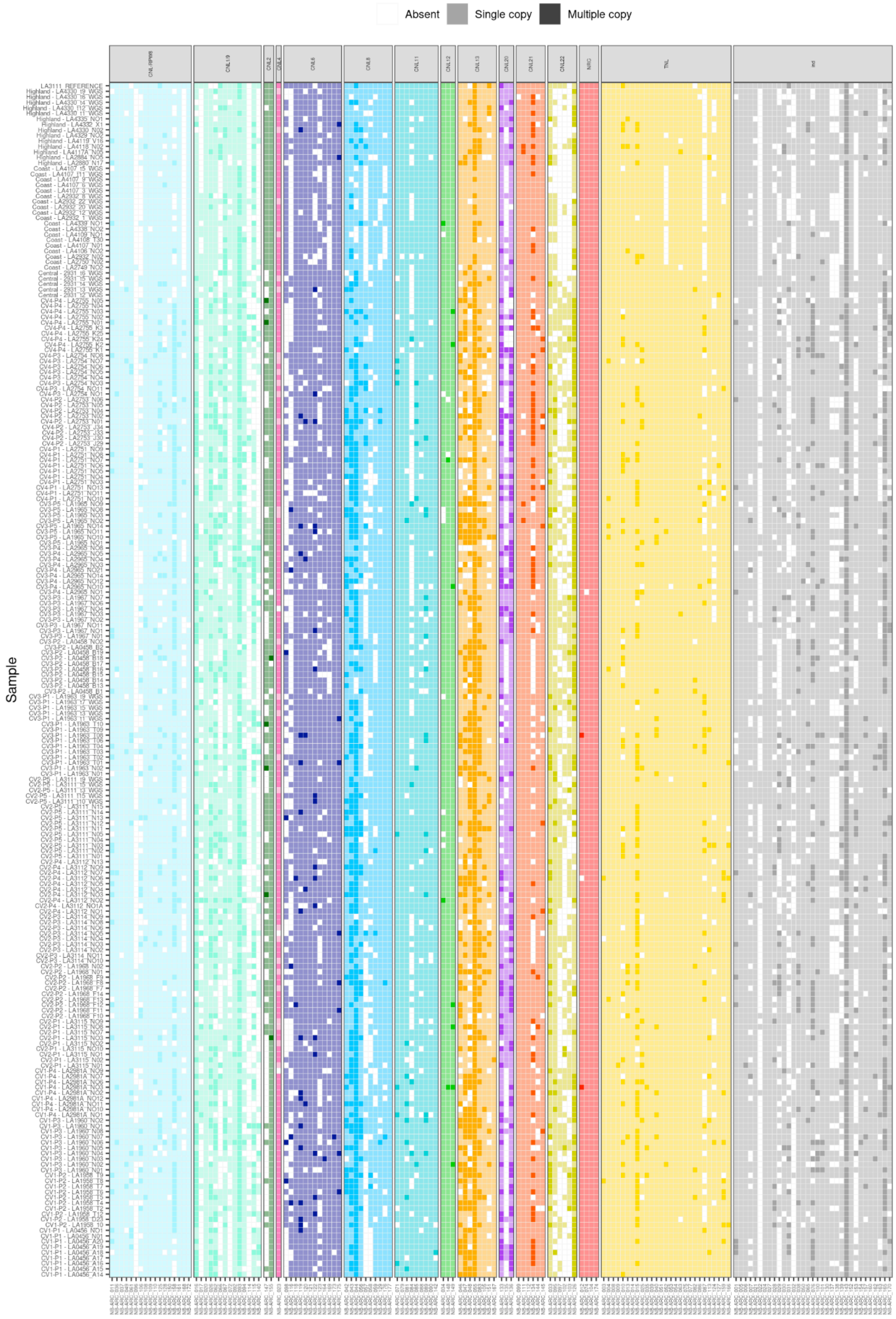

New Phytologist

**Figure S1.** Presence/Absence variation and copy number variation matrix of NB-ARC loci of *Solanum chilense* detected through *de novo* assembly approach across 186 samples included in the sequence capture data set, whole-genome data from the reference genome sample (LA3111_t13; Stam *et al.*, 2019) and 30 whole-genome sequencing approach from Wei *et al.* (2023). The loci/sample displayed in a darker color show multiple hits above the identity and coverage thresholds (see Figure S4) indicating possible CNV of the locus in each sample. This CNV signal is consistent with higher mapping rates than expected based on the mapping rates in other loci (see Figure S5).

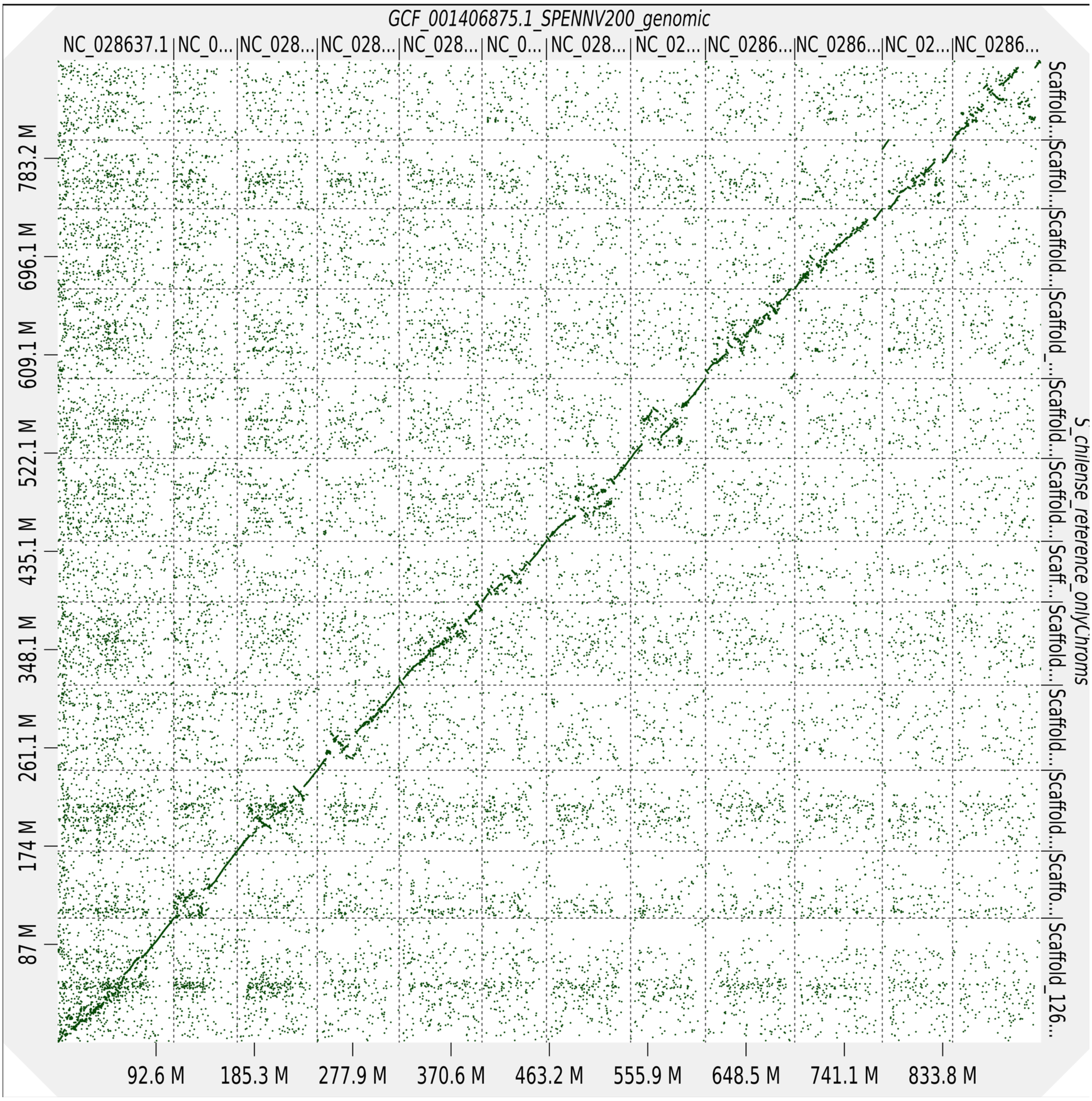

**Figure S2**. Dotplot representation of sequence alignment of the new assembly of *Solanum chilense* and *S. pennellii*.

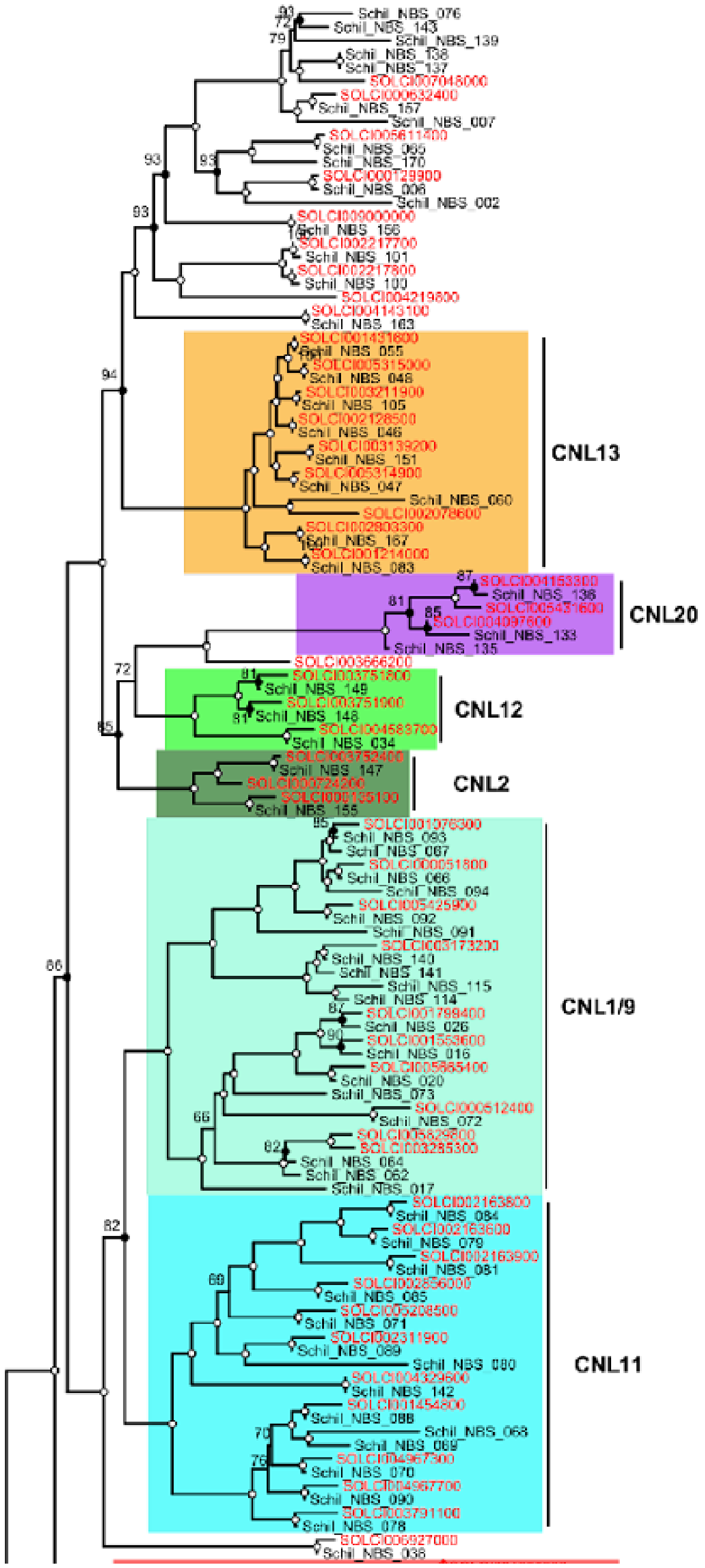
CNL13
CNL20
CNL12
CNL2
CNL1/9
CNL11

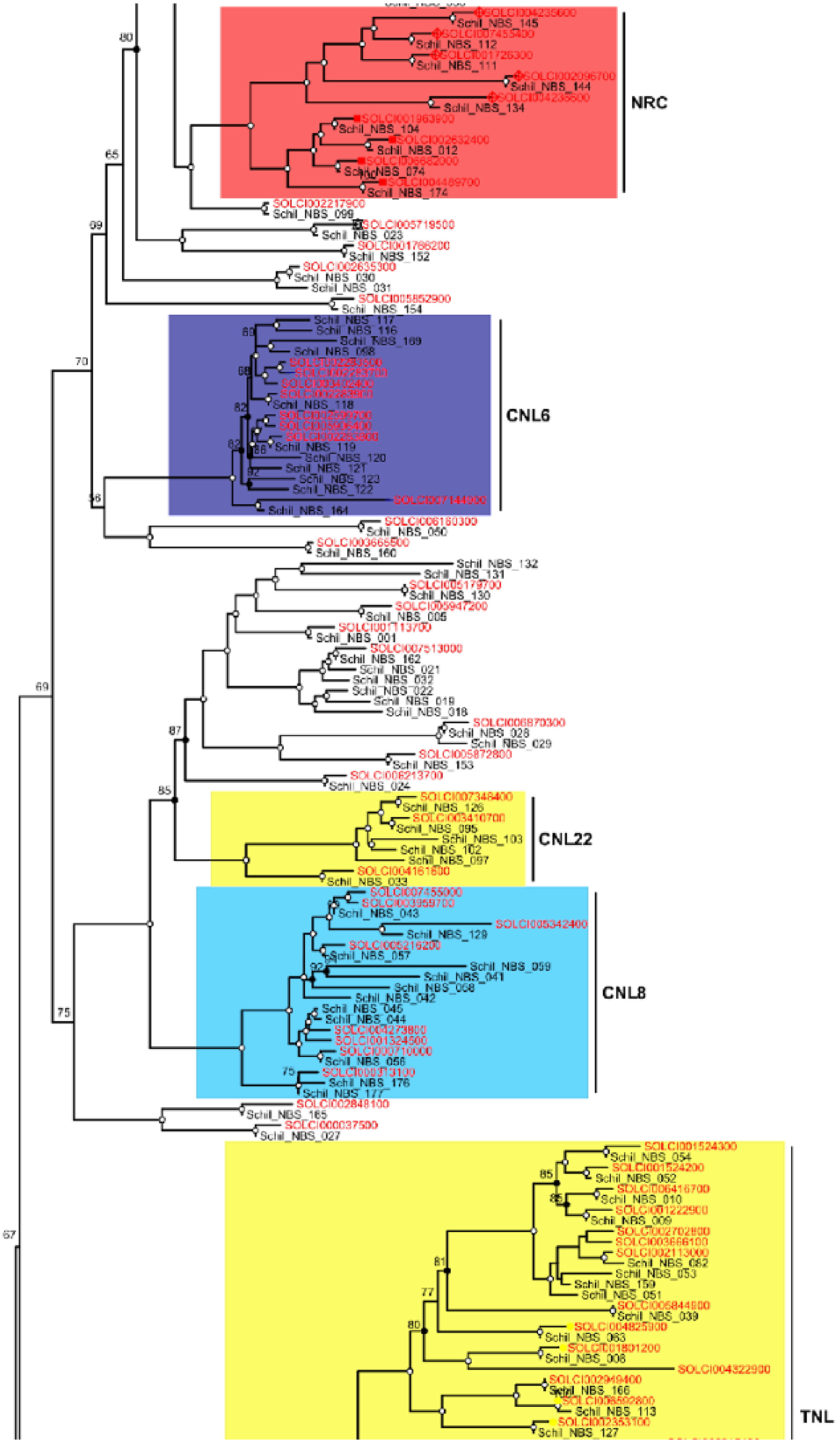
NRC
CNL6
CNL22
CNL8
TNL

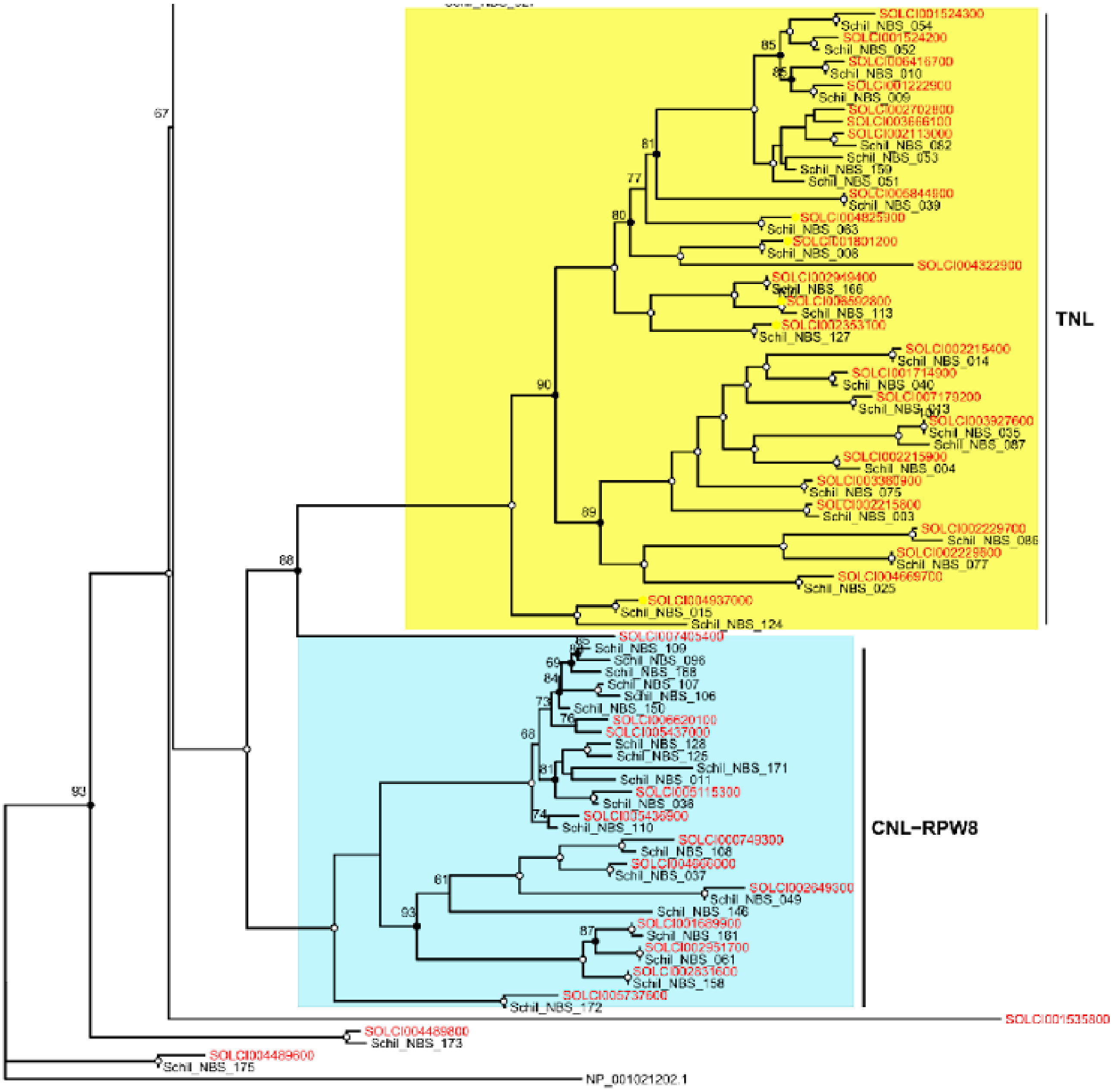


**Figure S3**. Maximum Likelihood phylogeny of the NB-ARC domains recovered from the Dovetail and the short-read sequence assembly from *S. chilense*, using NLR-PARSER. Clades are coloured according to the same colour scheme as used in Stam *et al.* (2019). Nodes with UltraFastBootstrap support of 95% and above are indicated with white circles, whereas branch support 80% to 94% are indicated with black circles. Numbers next to the left of the nodes indicate ultrafast bootstrap support, although these are not represented for nodes with support of 95% and above. Tip names in red represent NB-ARC sequences identified in Stam *et al.* (2019).

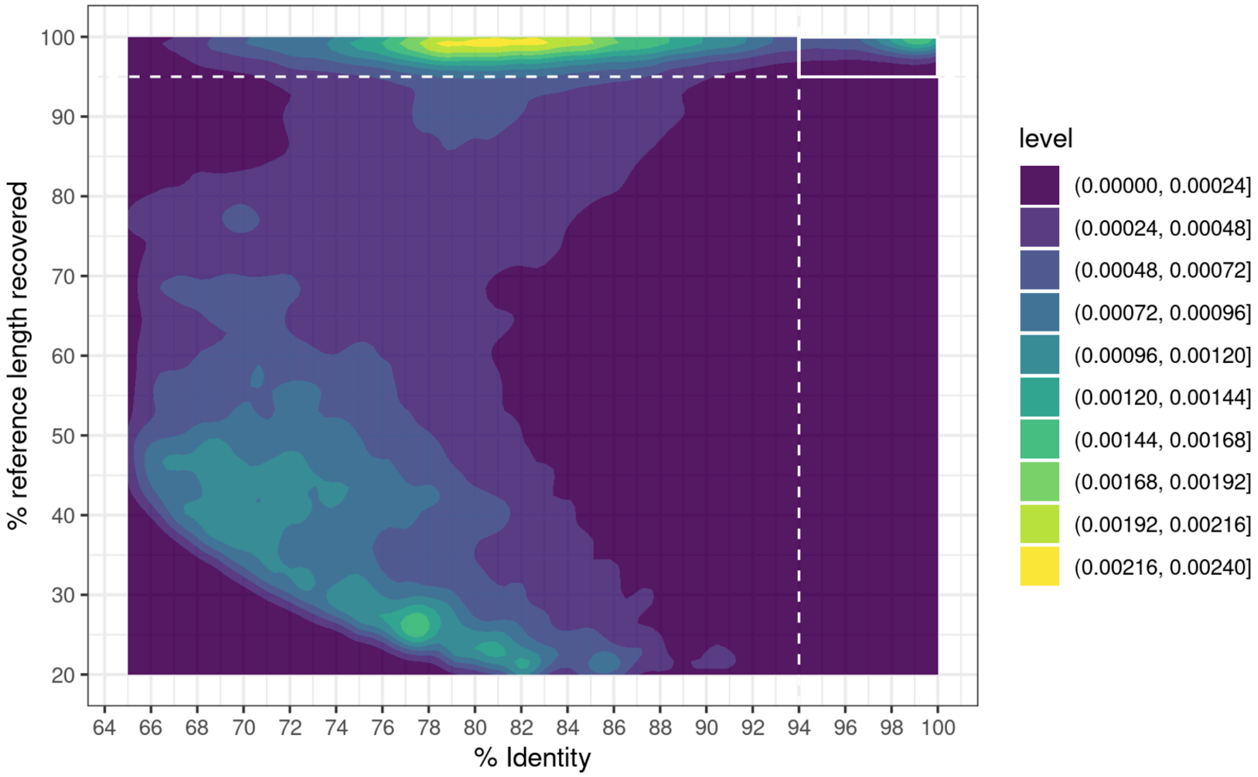


**Figure S4.** 2D density plot summarizing the identity percentage and the coverage percentage of all assembled contigs to the NB-ARC reference protein sequences. Dashed lines indicate the threshold values chosen to filter the extraction of *de novo* assembled loci in all samples.

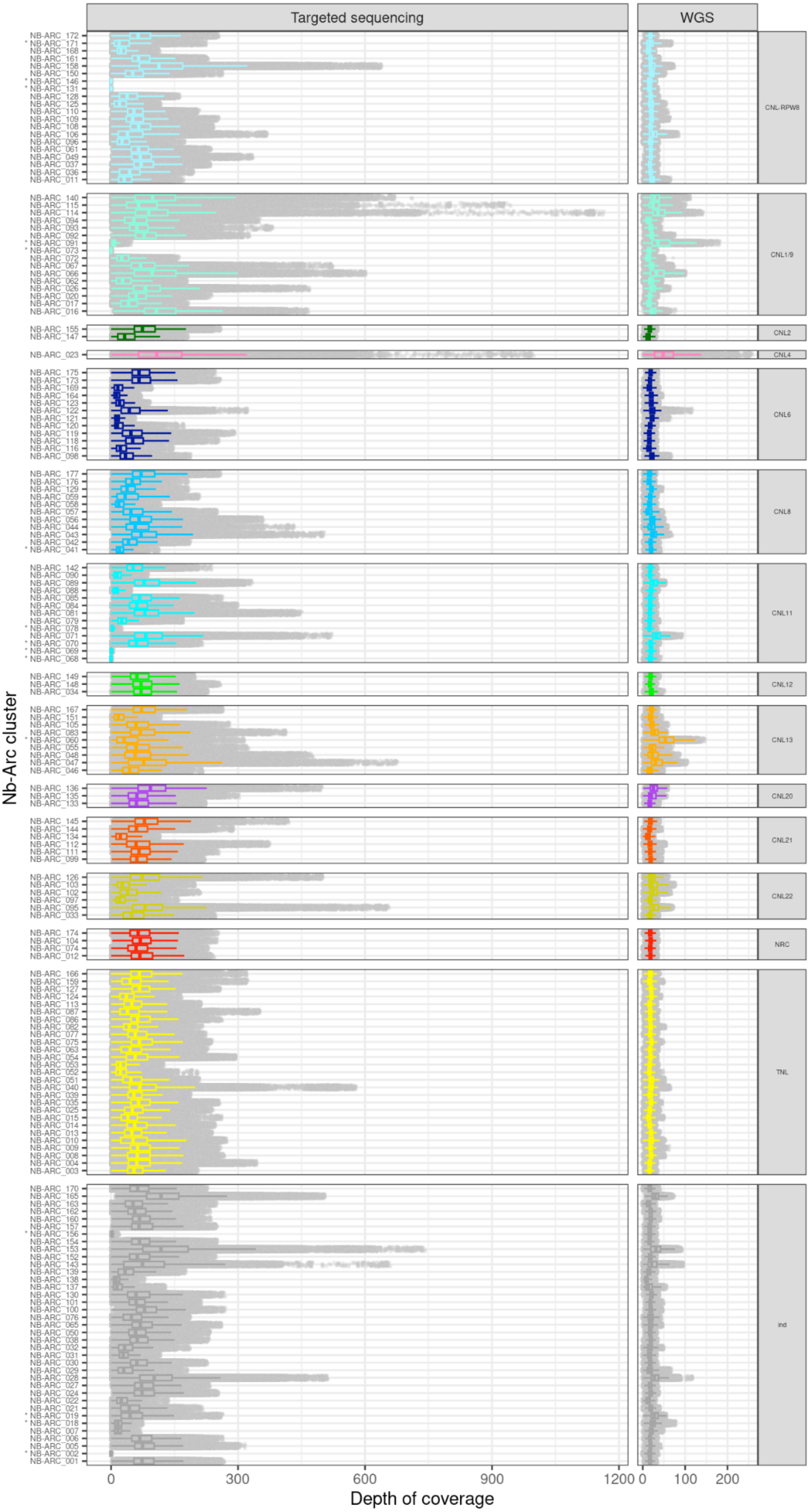
Targeted sequencing
WGS
Nb-Arc cluster
CNL-RPW8
CNL1/9
CNL2
CNL4
CNL6
CNL8
CNL11
CNL12
CNL13
CNL20
CNL21
CNL22
NRC
TNL
ind
0
300
600
900
1200
0
100
200
Depth of coverage

**Figure S5**. Distribution of depth of coverage for each NBS locus annotated in the new reference genome of *Solanum chilense* measured after read alignment of the 200 samples from the targeted sequencing (left panels) and 30 samples from whole-genome sequencing (right panels). NBS marked with an asterisk indicate removed loci.

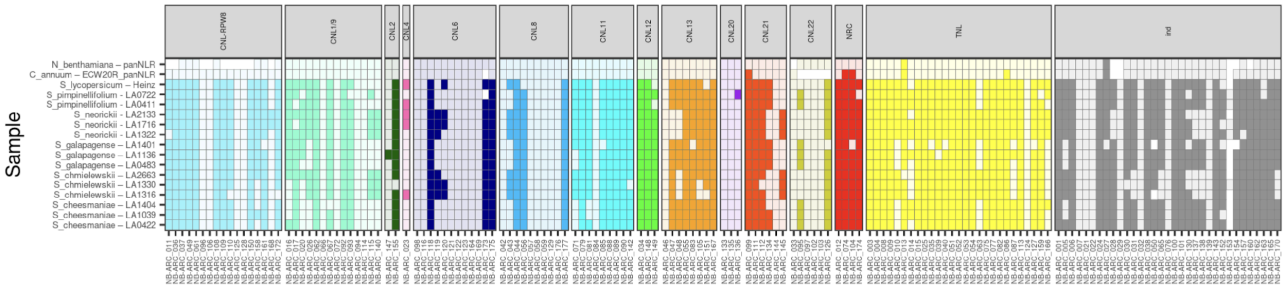


**Figure S6**. NB-ARC loci presence-absence variation matrix detected through *de novo* assembly approach for outgroup RENseq samples from Seong *et al.* (2020) that includes *S. lycopersicum*, five wild tomato species, and two additional Solanaceae.

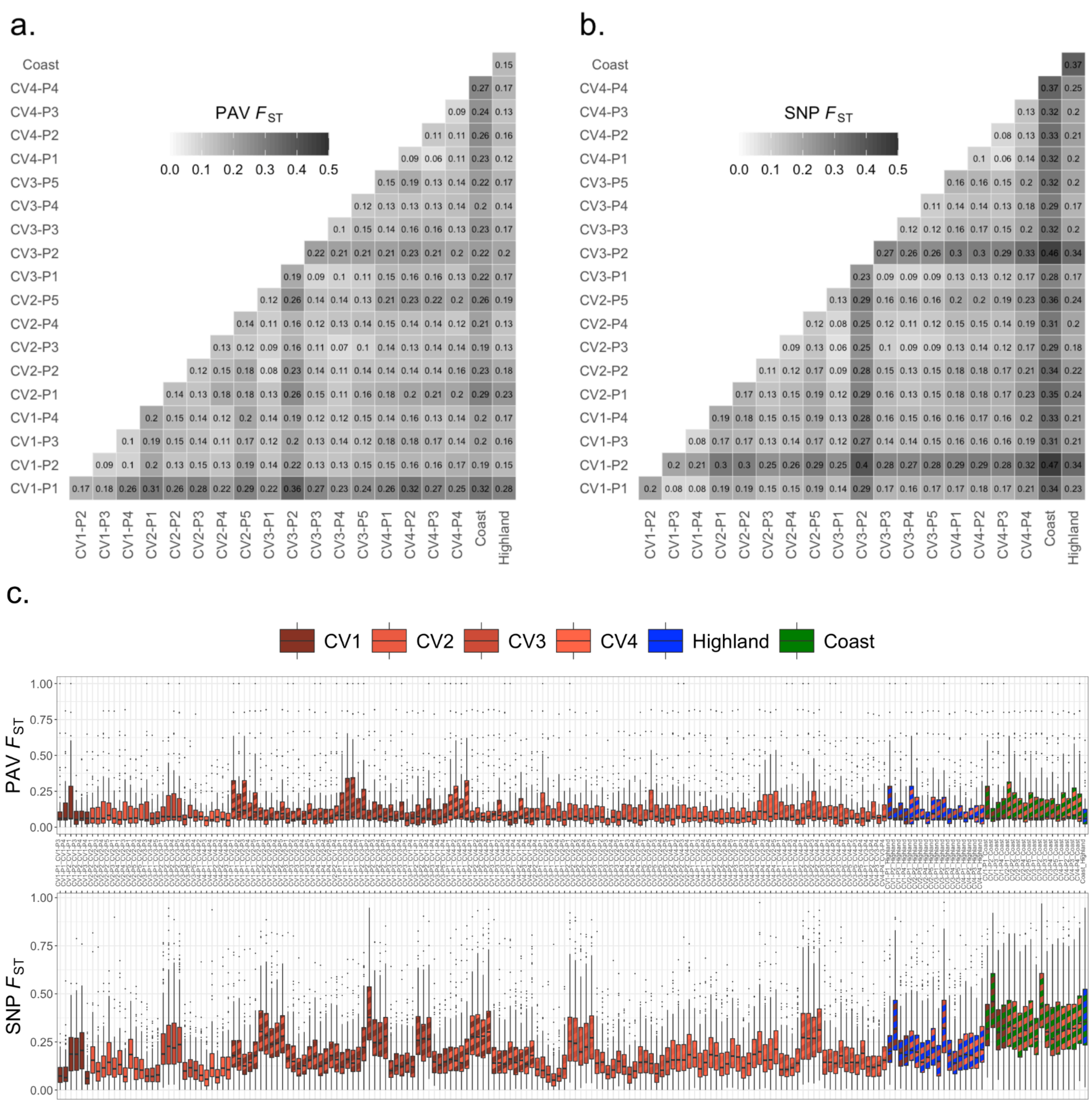


**Figure S7**: Population differentiation $F_{ST}$ estimated on NB-ARC (**a**) presence-absence variation (PAV) and (**b**) nucleotide variation (SNP). **c.** Distribution of airwise $F_{ST}$ distribution for PAV (top) and SNP (bottom).

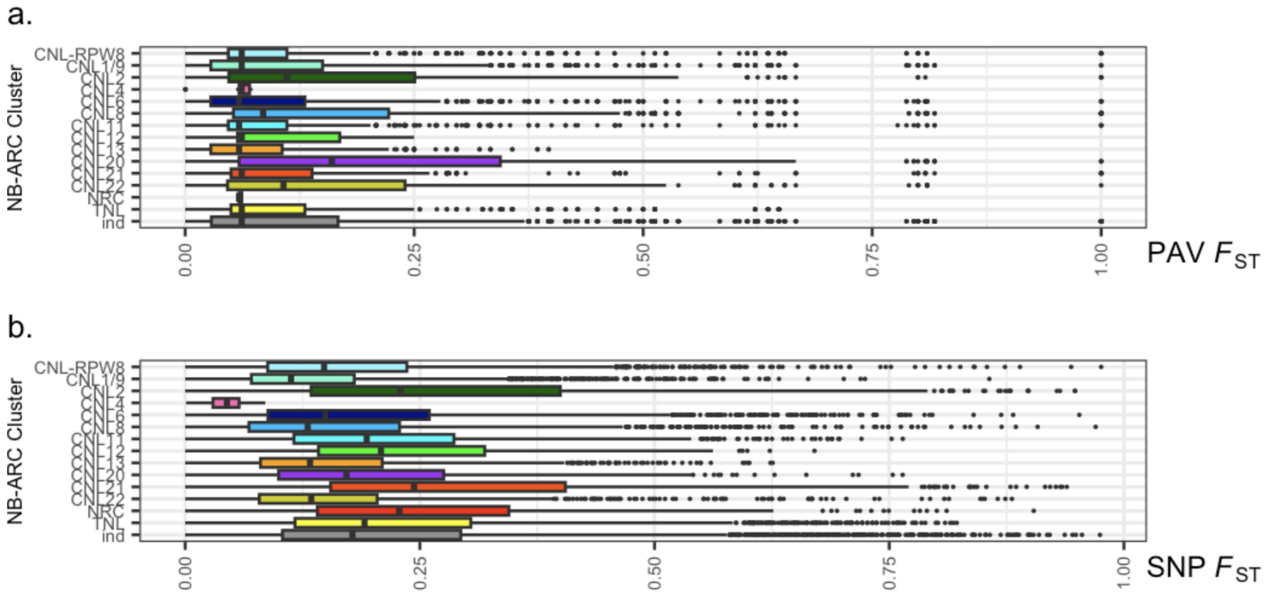


**Figure S8**: Distributions of per-locus NB-ARC pairwise population differentiation $F_{ST}$ estimated on presence-absence variation (PAV) and nucleotide variation (SNP) **a.** $F_{ST}$ distribution for PAV by NB-ARC cluster. **b.** $F_{ST}$ distribution for SNP data by NB-ARC cluster.

New Phytologist

#### Supplementary tables

**Table S1.** Comparison of the 170 NB-ARC-containing NLRs recovered in the new Dovetail genome versus the previous draft *S. chilense* genome (Stam *et al.*, 2019) for each NLR clades. Numbers in parentheses indicate the amount of new sequences identified in the DT genome.

<table>
<tr><th>NLR Clades</th><th>Helper/Sensor /Non-Sensor</th><th>Nb of sequences in DT genome</th><th>Nb of sequences in the G3 genome</th></tr>
<tr><td>CNL1/9</td><td>sensor</td><td>16 (8)</td><td>10</td></tr>
<tr><td>CNL11</td><td>sensor</td><td>13 (2)</td><td>11</td></tr>
<tr><td>CNL12</td><td>sensor</td><td>3</td><td>3</td></tr>
<tr><td>CNL13</td><td>sensor</td><td>9</td><td>9</td></tr>
<tr><td>CNL2</td><td>sensor</td><td>2</td><td>3</td></tr>
<tr><td>CNL20</td><td>sensor</td><td>3</td><td>3</td></tr>
<tr><td>CNL21</td><td>Helper (extended)</td><td>6 (1)</td><td>5</td></tr>
<tr><td>NRC</td><td>Helper</td><td>4</td><td>4</td></tr>
<tr><td>CNL22</td><td>Non-sensor</td><td>6 (3)</td><td>3</td></tr>
<tr><td>CNL6</td><td>Non-sensor</td><td>12 (7)</td><td>8</td></tr>
<tr><td>CNL8</td><td>Non-sensor</td><td>11 (8)</td><td>7</td></tr>
<tr><td>CNL-RPW8 (RNL)</td><td>Non-sensor</td><td>20 (11)</td><td>9</td></tr>
<tr><td>CNL4</td><td>Non-sensor</td><td>1</td><td>1</td></tr>
<tr><td>TNL</td><td>Non-sensor</td><td>27(5)</td><td>28</td></tr>
<tr><td rowspan="2">indetermined</td><td>Non-sensor</td><td>21 (7)</td><td rowspan="2">30</td></tr>
<tr><td>Sensor</td><td>16 (8)</td></tr>
</table>
